# Observation of Higher-Order Assemblies Controlled by Protein One-dimensional Movement

**DOI:** 10.1101/2024.06.06.597732

**Authors:** Xiao-Peng Han, Ming Rao, Shu-Rui Wu, Qiurong Zhang, Fajian Hou, Shao-Qing Zhang, Ling-Ling Chen, Jiaquan Liu

## Abstract

RIG-I-like receptors (RLRs) detect cytosolic viral RNAs and initiate innate immune response. MDA5 is an RLR that recognizes viral long double-stranded RNA (dsRNA). The central helicase and C-terminal domains of MDA5 form a ring-like clamp around dsRNA, and its activation requires the oligomerization of N-terminal caspase activation and recruitment domains (CARDs). Another RLR, LGP2, lacks CARDs but plays essential roles in MDA5 signaling. However, the mechanisms underlying MDA5 CARDs assembly and the collaborative mechanics between MDA5 and LGP2 remain largely unresolved. Here we found the MDA5 ring-like clamp operates as an ATP-hydrolysis-driven motor, with the self-assembly of its CARDs strictly suppressed by the protein one-dimensional (1D) translocation along dsRNA. LGP2 recognizes a mobile MDA5 and inhibits the motor’s movement, thereby allowing MDA5 CARDs to assemble into oligomers that can further interact with downstream components. These findings highlight how RLR sensors orchestrate higher-order assemblies through 1D movement to direct immune response.

## INTRODUCTION

The innate immune system is the first line of host defense against microbial pathogens and relies on pattern-recognition receptors (PRRs) that identify pathogen-associated molecular patterns (PAMPs)^1,2^. Among these PRRs, RIG-I-like receptors (RLRs) are cytosolic viral RNA sensors found in mammals, including RIG-I, MDA5, and LGP2. All three of these human RLRs share a highly similar structure, containing a DExD/H-box helicase domain and a CTD that both are crucial for RNA recognition^1–3^. RIG-I and MDA5 have two N-terminal CARDs that can mediate downstream signal transduction, with intrinsically disordered regions (IDRs, ~ 50 - 100 residues) connecting the CARDs to the helicase domains. LGP2 lacks the CARDs and is generally believed to regulate RIG-I and MDA5^4^.

The antiviral signaling begins with viral RNA recognition by the RLR sensors. Short dsRNA with a triphosphate group at the 5’ blunt end (5’-ppp) serves as an agonist for RIG-I, while MDA5 prefers long dsRNA fragments (> 1 kb) such as picornavirus replicative-form dsRNA^5^. The RNAs that act as ligands for LGP2 are poorly understood. Signaling downstream of RLRs requires the adaptor protein mitochondrial antiviral signaling protein (MAVS, also known as IPS-1, CARDIF, and VISA)^6–9^, which is localized to the mitochondrial outer membrane and comprises a CARD domain at its N terminus. When RIG-I or MDA5 is activated by viral RNAs, CARDs from proximate RLR proteins assemble into tetramers or higher-order oligomers that directly interact with MAVS CARD, leading to the formation of MAVS filament^10,11^. The MAVS filament further recruits TRAF molecules and activates the downstream signaling pathway^12^.

The proximity-induced higher-order assemblies have been regarded as a fundamental principle governing immune signaling^13^. To maintain cellular immune homeostasis, it is critical to suppress the spontaneous assembly of CARDs within RLRs^1,2^. Without viral RNA, RIG-I maintains in an auto-repressed state where the tandem CARDs associate with the helicase domain^14,15^. This configuration serves as a physical barrier to prevent the CARDs from forming tetramers, thus blocking signaling to MAVS^16,17^. Binding to viral dsRNA with a 5’-ppp induces a conformational rearrangement within RIG-I and releases the CARDs^14,15^. In contrast, MDA5 lacks the intrinsic CARDs-helicase interactions^11,17^; therefore, the progression of how RNA recognition leads to CARDs assembly is less well understood in the MDA5 pathway^1,2^. Early reports have shown that the helicase and CTD domains of MDA5 formed a ring-like clamp around dsRNA^18^. Biochemical studies suggested multiple MDA5 rings were arranged head-to-tail along the dsRNA axis, potentially creating a filamentous, highly organized structure^19–22^. It was further hypothesized that the assembly of an MDA5 filament occurred specifically on long dsRNA molecules, leading to the oligomerization of CARDs in close proximity, thereby activating MAVS^23,24^. LGP2 and the ATP cycle of MDA5 were thought to regulate the MDA5 filament formation, although their roles in signaling remain elusive^25,26^.

In recent decades, significant observations have emerged regarding proteins moving along nucleic acids backbone in a one-dimensional (1D) manner. These movements involve various mechanisms such as ATP-hydrolysis-driven translocation, passive sliding/diffusion, intersegmental transfer, or hopping/jumping^27,28^. Additional evidence indicates extensive utilization of 1D motions by DNA-binding proteins during DNA repair, DNA replication, and genome maintenance. Their biological functions are often distance-related, for example, facilitating target search^29,30^, enhancing processivity^31,32^, enabling long-range communications^33^, and driving loop extrusion^34–36^. A few RNA-binding proteins, including RIG-I, have been reported to translocate on RNA^37^. Further investigations suggested that this 1D movement might not directly contribute to signaling, leaving its biological significance largely unknown^38^. Nonetheless, despite the considerable focus on these biophysical behaviors themselves, little attention has been paid to their potential influence on protein higher-order assemblies.

Here, we employed single-molecule imaging to investigate MDA5 signaling in real-time. We reveal that the helicase and CTD of MDA5 function together as an ATP-dependent motor that translocates along dsRNA. The 1D movements readily induce CARDs-CARDs interactions, facilitating the cooperative loading of multiple MDA5 motors onto a RNA substrate. However, these MDA5 motors, when moving in close proximity, are incapable of forming functional oligomers. The assembly of CARDs tetramers only occurs when the 1D motions have been completely impeded by LGP2 binding or by a non-hydrolysable ATP-analog. Our findings provide insights into how immune sensors control signaling activity by utilizing protein 1D movement.

## RESULTS

### MDA5 is a dsRNA motor powered by ATP

We exploited prism-based single-molecule total internal reflection fluorescence (smTIRF) microscopy^39,40^ to visualize individual MDA5 molecules on dsRNA. Single 11.6-kb dsRNA molecules were constructed by *in vitro* transcription and stretched across a passivated custom-made flow cell surface by laminar flow and linked at both ends via biotin-neutravidin interactions (Figures 1A, 1B, S1A and S1B; Table S1). Human MDA5 protein was purified and labeled with a Cy3 fluorophore similar to our previous studies^40^ (Figures S1C and S1D; Table S2). Injection of Cy3-MDA5 with ATP into the flow cell resulted in numerous single particles exhibiting one-dimensional (1D) movement along the dsRNA (Figure 1C, + ATP; Figures S2A and S2B; Movie S1). Interestingly, nearly all of MDA5 molecules moved unidirectionally without a reversal of direction, resembling the motions of a mechanochemical motor or an enzyme translocating along a double-stranded DNA (dsDNA) molecule^30,41,42^ (Figures 1C, S2A and S2B). The translocation of MDA5 was completely abolished in conditions lacking ATP or when ATP was substituted by a non-hydrolysable ATP-analog adenosyl-methylene-triphosphate (ADPCP) (Figures 1C and 1D, −ATP and +ADPCP), demonstrating that ATP hydrolysis is required for the 1D movement along dsRNA. In alignment with the single-molecule analysis, ATPase activity of MDA5 was stimulated upon the addition of 11.6-kb dsRNA (k_cat_ = 1.1 ± 0.1 s^−1^), but not ssRNA (Figures S2C and S2D). These observations directly show that individual MDA5 molecule operates as a motor, translocating along dsRNA using the energy derived from ATP hydrolysis.

**Figure 1.**
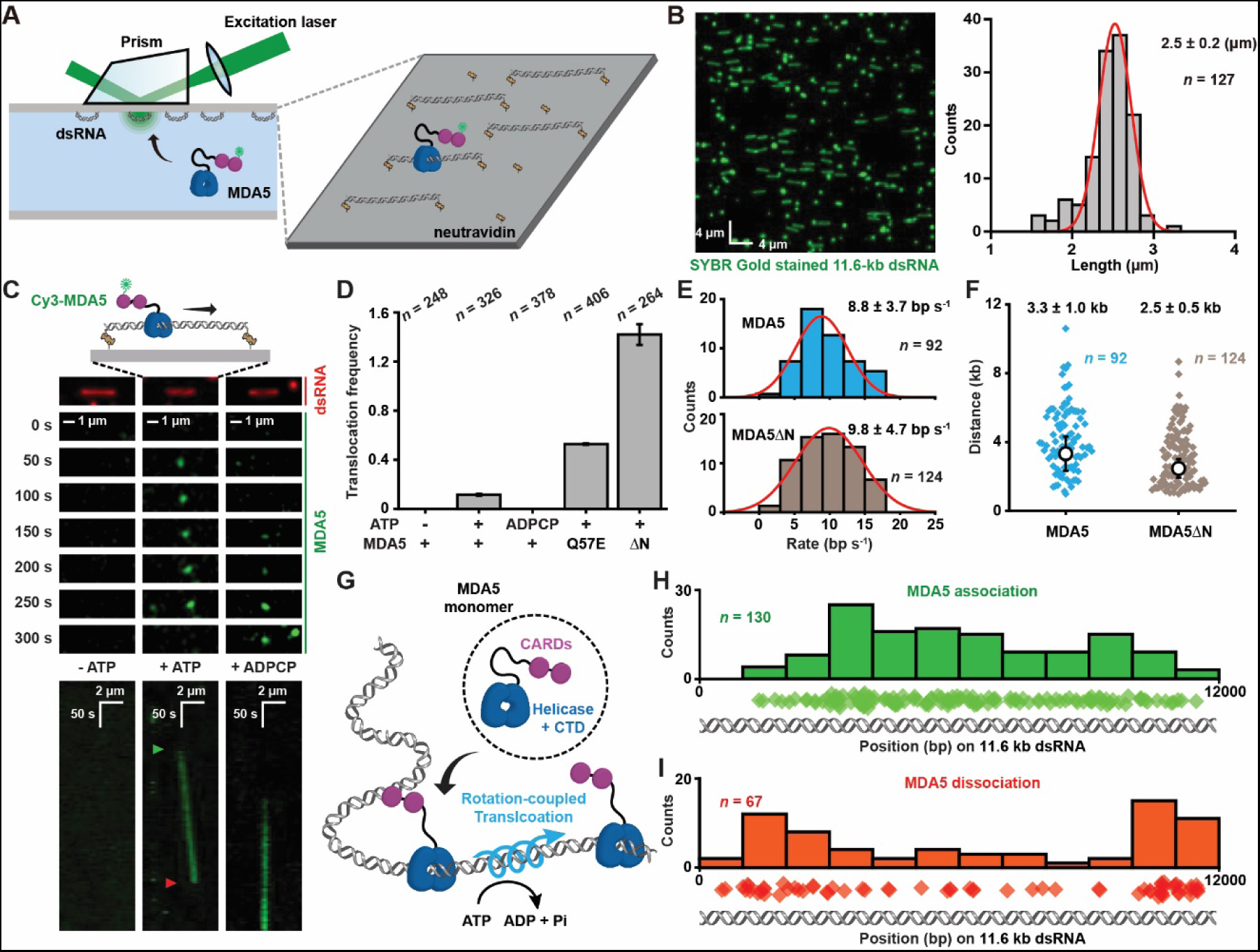
MDA5 is an ATP-hydrolysis-driven motor on dsRNA. (A) A schematic illustration of dsRNA and MDA5 observations by prism-based smTIRF microscopy. (B) Left: representative 11.6-kb dsRNA visualized by smTIRF microscopy in the absence of flow. The dsRNA was stained with SYBR Gold and a 42.7 x 42.7 µm field of view is shown. Right: the length distribution of the 11.6-kb dsRNA observed by smTIRF microscopy (*n* = number of dsRNA molecules). The data was fit with a Gaussian distribution that determined the mean ± s.d.. The contour length of an 11.6-kb dsRNA is calculated to be 3.3 µm. (C) Top: representative fluorescent images and a schematic illustration showing the translocation of a Cy3-MDA5 motor on 11.6-kb dsRNA with ATP (middle, + ATP), and the absence of any MDA5 motors without ATP or with ADPCP (left and right, −ATP and + ADPCP). SYBR Gold stained 11.6-kb dsRNA is shown in red and Cy3-MDA5 at various times is shown in green. Bottom: representative kymographs of Cy3-MDA5 on dsRNA under various conditions. Kymographs were generated by images stacking (see Figure S2A). Green and orange arrowheads indicate the association and dissociation of MDA5 motor on dsRNA, respectively. (D) The frequency of MDA5, MDA5(Q57E) or MDA5ΔN motor (3 nM) translocation on 11.6-kb dsRNA under various conditions observed by smTIRF (mean ± s.d.; *n* = number of dsRNA molecules). (E) Histogram of binned MDA5 (top) and MDA5ΔN (bottom) translocation rates that were fit to Gaussian function to derive the average rates (mean ± s.d.; *n* = number of events). (F) Distribution of MDA5 and MDA5ΔN translocation distance. Diamonds represent individual events and open circles represent the mean processivity derived by single exponential decay fitting (mean ± s.e.; *n* = number of events). (G) A schematic illustration showing the rotation-coupled translocation of a single MDA5 molecule along dsRNA backbone. (H) Distribution of the starting positions for MDA5 translocation on dsRNA (*n* = number of events). Diamonds represent individual starting events. (I) Distribution of the ending positions for MDA5 translocation on dsRNA (*n* = number of events). Diamonds represent individual ending events.

While the helicase domain and CTD of MDA5 are responsible for substrate recognition^18^, it remains unclear whether the CARDs participate in the 1D translocation. A mutant MDA5(Q57E), which had impaired CARDs-CARDs interactions^11^, and a CARDs-deleted MDA5 (MDA5ΔN) proteins were purified, labeled and introduced into smTIRF (Figures S1C and S1D). Both the mutant and truncated proteins largely retained the dsRNA-dependent ATPase activities (Figure S2D). Remarkably, the frequencies of MDA5(Q57E) and MDA5ΔN translocation events were ~5-fold and 12-fold higher, respectively, compared to wild-type MDA5, whereas the translocation rates and processivities between MDA5 and MDA5ΔN appeared nearly identical (Figures 1D-1F and S2E). These findings suggest MDA5 motor always initiates translocation as a monomer, and it is likely that the CARDs-CARDs interactions occur between MDA5 molecules, leading to a decrease in monomeric protein concentration for wild-type MDA5 (Figure 1G). Notably, the translocation speed of MDA5 was relatively slow (~ 9 bp s^−1^) compared with many other DNA translocases^30,41^, indicating that the motor maintained tight contact with the RNA phosphate backbone^18^ (Figure 1E). Our data, along with the observations of MDA5 ring-like clamps twisting along the dsRNA backbone in structural analyses^22,43^, strongly implies that the 1D movement is rotation-coupled, a mechanism enabling the motor to closely track an RNA double helix (Figure 1G).

The initial binding of the MDA5 motor to its substrate appeared to occur randomly across the entire length of dsRNA (Figures 1C and 1H), indicating that MDA5 recognized internal dsRNA backbone without dependence on specific sequences. In contrast, most of the dissociation positions of MDA5 were either at or very near the ends of dsRNA, suggesting the motors were released upon encountering an RNA end (Figures 1C and 1I). We purified Cy5-labeled RIG-I (Figures S1C and S1D), then examined the interactions between RIG-I and an 11.6-kb dsRNA anchored at a single end (Figure S3A). In the presence of ATP, the majority of RIG-I molecules were found to bind to locations near the free end of dsRNA that contains a 5-’ppp, and no processive translocation event was observed (Figures S3B and S3C). These results are consistent with the notion that RIG-I mainly identifies the 5’-ppp ends of dsRNA substrates, while MDA5 recognizes internal regions within long dsRNA substrates^1^.

### LGP2 binding turns MDA5 motor off

The predicted structure of the human LGP2 is similar to that of an MDA5 motor lacking CARDs (Figure 2A). We found LGP2 also maintained a dsRNA-dependent ATPase activity (Figure 2B), although the rate of ATP hydrolysis by LGP2 was ~ 20 times slower than that of MDA5 (k_cat_ = 0.062 ± 0.002 s^−1^; compare Figures S2D with 2B). However, it remains uncertain whether LGP2 functions as an ATP-dependent motor on dsRNA. To explore the interactions between LGP2 and 11.6-kb dsRNA, we purified recombinant LGP2 and labeled it with a fluorophore (Figures S1C and S1D). Upon injecting Cy3-labeled LGP2 into the flow cell without ATP or MDA5, we observed numerous LGP2 molecules localizing to the internal sites within the double-end-tethered dsRNA, resulting in the formation of a string-like dsRNA molecule coated with fluorescent LGP2 (Figures 2C and 2D). Addition of ATP eliminated most of LGP2-dsRNA interactions, with no observable LGP2 translocation events (Figures 2C and 2D). Interestingly, co-injection of LGP2 and MDA5 with ATP retained most LGP2 on the dsRNA, leading to the formation of numerous LGP2-coated dsRNA (Figures 2C and 2D). To further determine the roles of ATP in LGP2-dsRNA interactions, an ATP-binding deficient mutant LGP2(K30A)^4,44^ was purified, labeled (Figures S1C and S1D), and shown to lack dsRNA-dependent ATPase activity^4,44^ (Figure 2B). As expected, LGP2(K30A)-dsRNA interactions remained unaffected by the addition of ATP, resulting in LGP2(K30A)-coated dsRNA under all conditions (Figures 2C and 2E). These findings suggest that LGP2 functions differently from an MDA5 ring-like clamp, and that the principal role of ATP is to release LGP2 from internal sites of dsRNA (Figure 2F). Given the abundance of cellular ATP molecules, it is likely that MDA5 is required to load LGP2 on long dsRNA substrates (Figure 2F).

**Figure 2.**
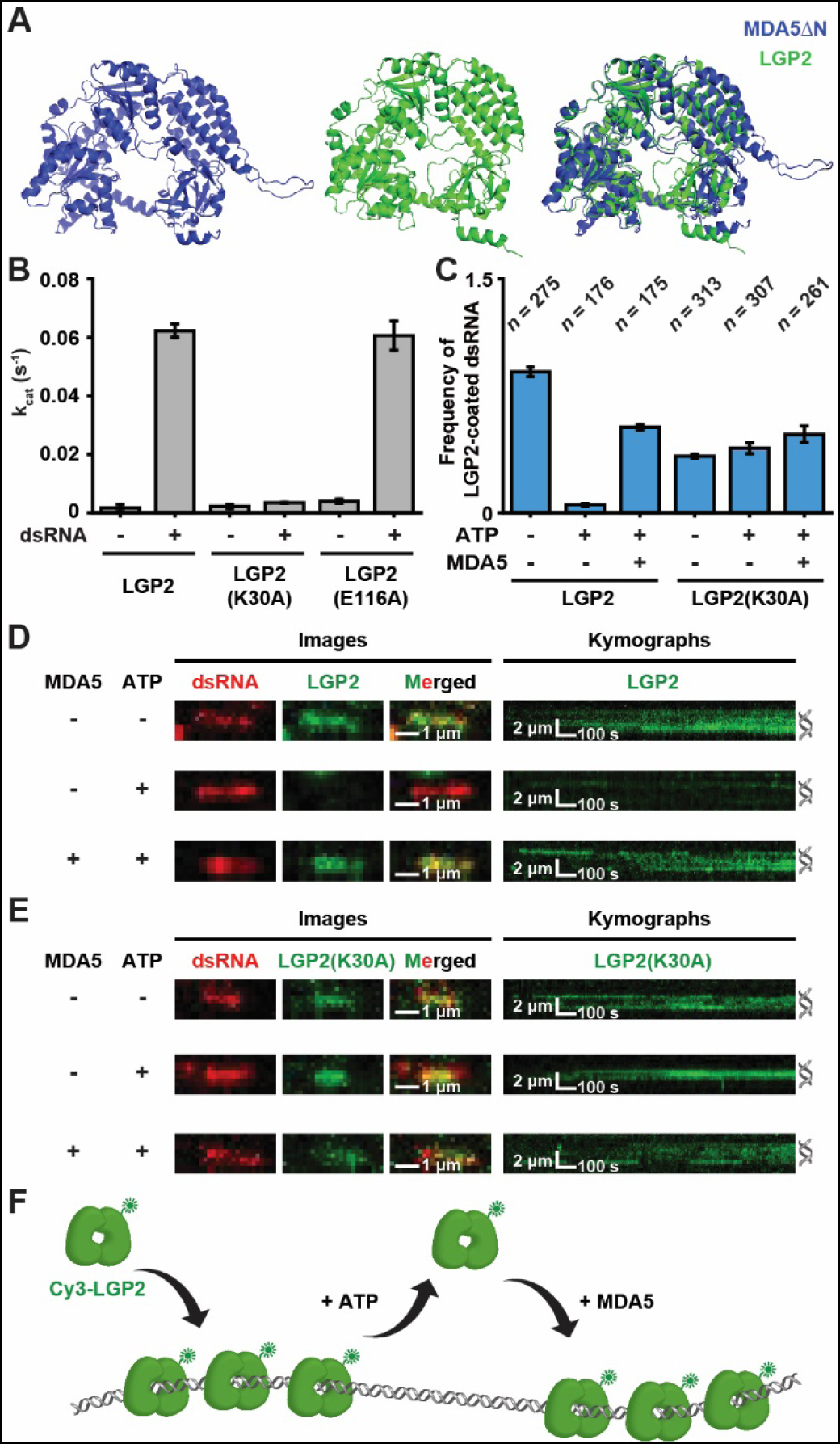
ATP recycles LGP2 from long dsRNA. (A) AlphaFold predicted structures of human MDA5ΔN (left), LGP2 (middle) and the pairwise structure alignment of two proteins (right). (B) The turnover numbers (k_cat_) of LGP2, LGP2(K30A) and LGP2(E116A) ATPase using 11.6-kb dsRNA substrate (error bars: mean ± s.e.). (C) The frequency of LGP2-coated dsRNA under various conditions (mean ± s.d.; *n* = number of dsRNA molecules). (D) Representative images (left) and kymographs (right) showing the binding of 60 nM Cy3-LGP2 on dsRNA under various conditions. SYBR Gold stained 11.6-kb dsRNA is shown in red and Cy3-LGP2 (20 min incubation after injection) is shown in green. Merged images are generated by overlaying both channels. Positions of dsRNA are shown adjacent to the right of kymographs. (E) Representative images (left) and kymographs (right) showing the binding of 60 nM Cy3-LGP2(K30A) on dsRNA under various conditions. SYBR Gold stained 11.6-kb dsRNA is shown in red and Cy3-LGP2(K30A) (20 min incubation after injection) is shown in green. Merged images are generated by overlaying both channels. Positions of dsRNA are shown adjacent to the right of kymographs. (F) A schematic illustration showing that ATP dissociates Cy3-LGP2 from dsRNA and MDA5 recruits LGP2 onto dsRNA.

The recruitment of LGP2 through an MDA5-dependent mechanism underscores the existence of physical interactions between the two helicases (Figure 2). Single-molecule tracking revealed that co-injection of both proteins resulted in frequent co-localization of Cy3-MDA5 and Cy5-LGP2 particles on the dsRNA (Figures 3A, 3B and S4A; Movie S2). Remarkably, the binding by LGP2 appeared to stop MDA5 translocation, leading to the formation of an immobile MDA5-LGP2 complex at internal sites of the dsRNA (Figures 3A-3C and S4A; Movie S2). Immobile complexes were still observed when wild-type MDA5 was substituted by MDA5ΔN (Figure S4B), indicating the MDA5-LGP2 interactions are CARDs-independent. We modeled the interaction interfaces using AlphaFold3 and found that several residues, K856 and K894 in MDA5, as well as E116, E117 and D603 in LGP2, seemed to be important for the MDA5-LGP2 interactions (Figure 3D).

**Figure 3.**
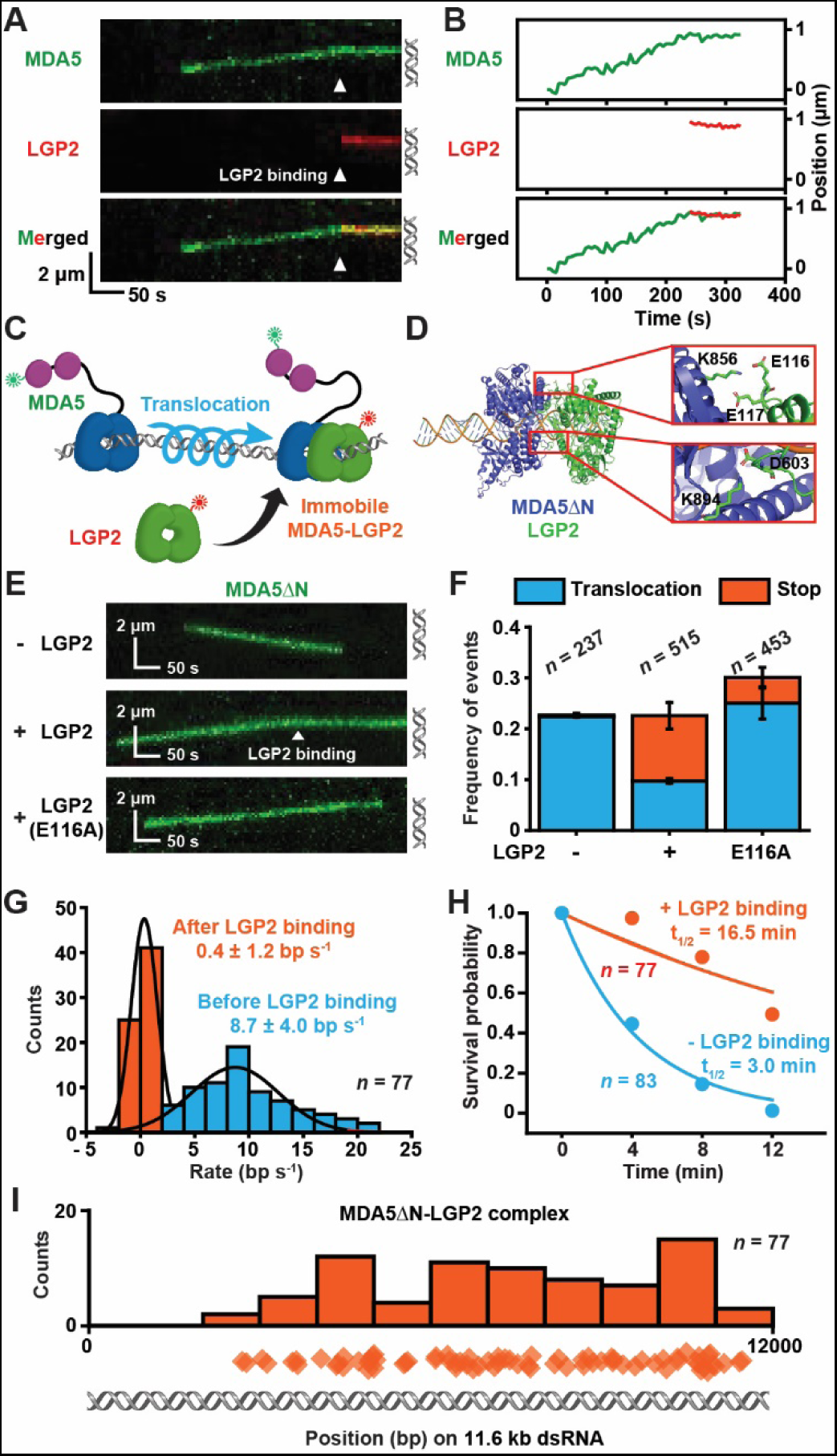
LGP2 binding stops MDA5 translocation on dsRNA. (A) Representative kymographs showing a Cy5-LGP2 (3 nM) binding to a Cy3-MDA5 motor (3 nM). Cy3-MDA5 is shown in green and Cy5-LGP2 is shown in red. Merged kymograph is generated by overlaying both channels. Arrowheads indicate the association of LGP2 with MDA5. Positions of dsRNA are shown adjacent to the right of kymographs. (B) Single-particle trajectories of MDA5 and LGP2 in Figure 3A. (C) A schematic illustration of LGP2 binding results in a stop in MDA5 translocation. (D) AlphaFold model of an MDA5ΔN-LGP2 complex on a 44-bp dsRNA substrate. The closeups show the key contacts between MDA5ΔN and LGP2. (E) Representative kymographs showing the translocation of a Cy3-MDA5ΔN motor (0.5 nM) on 11.6-kb dsRNA in the absence or presence of 0.5 nM LGP2/LGP2(E116A). Arrowheads indicate the association of LGP2 with MDA5ΔN. Positions of dsRNA are shown adjacent to the right of kymographs. (F) The frequency of MDA5ΔN translocation (blue) and LGP2-induced stop (orange) events under various conditions (mean ± s.d.; *n* = number of dsRNA molecules). (G) Histogram of binned MDA5ΔN translocation rates before and after LGP2 binding. Data were fit to Gaussian function to derive the average rates (mean ± s.d.; *n* = number of events). (H) Survival probability of MDA5ΔN and MDA5ΔN-LGP2 molecules on dsRNA. Data were fit by exponential decay functions to obtain a survival half-life (t_1/2_, mean ± s.e.; *n* = number of events examined). (I) Distribution of the positions for MDA5ΔN-LGP2 complex formation on dsRNA (*n* = number of events). Diamonds represent individual events.

A mutant LGP2 [LGP2(E116A)] was purified, shown to maintain dsRNA-dependent ATPase activity (Figure 2B), and introduced into smTIRF to validate the specific interactions between MDA5ΔN and LGP2 (Figures S1C and S1D). The translocation events of MDA5ΔN were found to be highly processive in the absence of LGP2, whereas the addition of LGP2 resulted in a stop in most of MDA5ΔN translocation events (~ 60%, Figures 3E and 3F). Substitution of wild-type LGP2 by LGP2(E116A) largely reduced the frequency of stop events (~ 17%, Figures 3E and 3F), suggesting that the single E→A mutation at interface could significantly diminish MDA5-LGP2 interactions. We calculated the translocation velocities of individual MDA5ΔN motor before and after LGP2 binding, and found that the MDA5ΔN-LGP2 interactions had completely shut the motor down on dsRNA (8.7 ± 4.0 bp s^−1^ before LGP2 binding, 0.4 ± 1.2 bp s^−1^ after LGP2 binding, Figure 3G). The MDA5ΔN motor alone typically dissociated from RNA ends, resulting in a short lifetime on RNA (-LGP2 binding, t_1/2_ = 3.0 min, Figure 3H). Conversely, once established, a stationary MDA5ΔN-LGP2 complex appeared to be stable and failed to reach the RNA ends, thereby extending its lifetime on dsRNA (+ LGP2 binding, t_1/2_ = 16.5 min, Figure 3H). The initial positions of MDA5ΔN-LGP2 complex formation were random across the length of a dsRNA (Figure 3I), which is consistent with the idea that LGP2 mainly recognizes a translocating MDA5 motor on a dsRNA. Collectively, these findings indicate that LGP2 binding acts as a “switch” for controlling the translocation activity of MDA5 along dsRNA.

### MDA5 motors form CARDs-dependent foci

Previous studies have shown that MDA5 proteins form a filament on dsRNA through helicase-helicase interactions, which further promotes CARDs oligomerization^19,23,25^. MDA5 protein concentrations were increased under smTIRF to allow multiple motors to be loaded onto a single dsRNA. We observed numerous wild-type MDA5 proteins forming foci while simultaneously translocating along dsRNA, with their assembly and disassembly being highly dynamic (Figures 4A, 4B and S4C; Movie S3). The fluorescent intensities of an MDA5 foci often increased more than 10-fold compared to that of a single protein, suggesting these foci contained at least a dozen MDA5 molecules. The foci were observed to form internally along the 11.6-kb dsRNA during protein translocation, and they immediately dissolved upon reaching a dsRNA end (Figures 4A and S4C; Movie S3). A fraction of MDA5 foci translocation (34 % of observed events; *n* = 91 / 269) resulted in the displacement of neutravidin from RNA ends, an ATP-dependent activity reported in a recent study^45^, leading to the withdrawal of the stretched dsRNA (Figure S4C, right). These results are consistent with previous observations that MDA5 proteins cooperatively bind to a long dsRNA substrate^19,20^.

**Figure 4.**
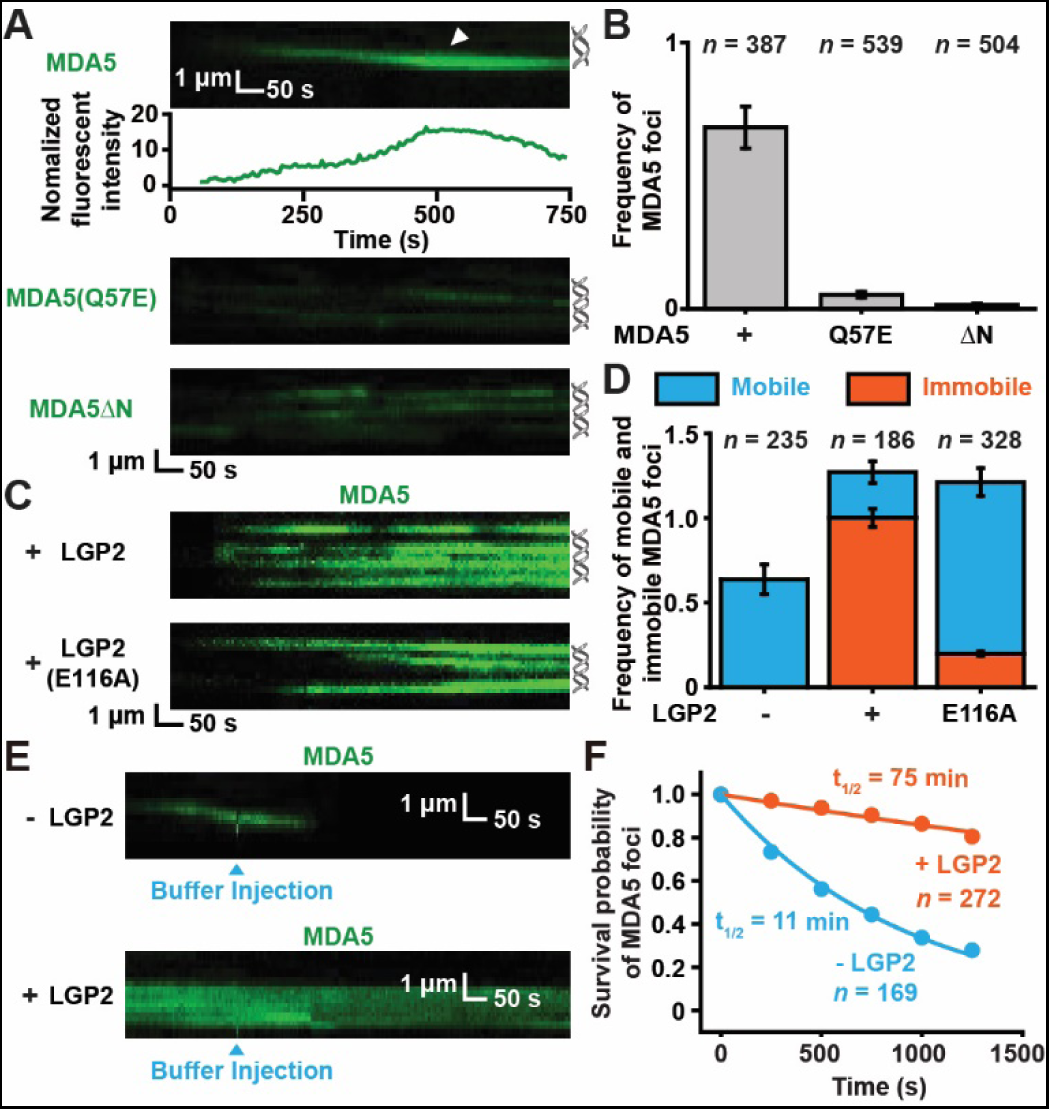
MDA5 motors form CARDs-dependent foci during 1D translocation. (A) Top: representative kymographs and time-dependent fluorescent intensities showing the cooperative binding of multiple Cy3-MDA5 motors (15 nM) to a dsRNA substrate. Arrowhead indicates the formation of a mobile MDA5 foci. Bottom: representative kymographs showing the absence of cooperative binding by Cy3-MDA5(Q57E) or Cy3-MDA5ΔN. Positions of dsRNA are shown adjacent to the right of kymographs. (B) The frequency of MDA5, MDA5(Q57E) or MDA5ΔN foci formation (mean ± s.d.; *n* = number of dsRNA molecules). (C) Representative kymographs showing the formation of Cy3-MDA5 foci in the presence of 10 nM LGP2/LGP2(E116A). Positions of dsRNA are shown adjacent to the right of kymographs. (D) The frequency of mobile (blue) and immobile (orange) MDA5 foci under various conditions (mean ± s.d.; *n* = number of dsRNA molecules). (E) Representative kymographs showing the stabilities of MDA5 foci and MDA5-LGP2 foci. Blue arrowheads indicate the injections of Imaging Buffer. (F) Survival probability of MDA5 foci and MDA5-LGP2 foci on dsRNA. Data were fit by exponential decay functions to obtain a survival half-life (t_1/2_, mean ± s.e.; *n* = number of events examined).

However, the vast majority of MDA5 foci were found to continuously move towards a single direction on dsRNA (Figures 4A, 4B and S4C; Movie S3), suggesting that MDA5 proteins within foci likely operated as multiple motors, rather than contributing to a bidirectionally elongating filament^23,25^. We replaced wild-type protein with MDA5(Q57E) or MDA5ΔN to determine the interactions that are essential for cooperative protein binding. Surprisingly, although numerous independent translocation events were observed on a single dsRNA, only a very small number of MDA5(Q57E) and MDA5ΔN foci were detected (Figures 4A and 4B). These data indicate that the formation of MDA5 foci requires CARDs-CARDs interactions, and it appears that the helicase-helicase contacts are insufficient for MDA5 oligomerization, at least under the protein concentrations used in smTIRF (15 nM, Figures 4A and 4B). Taken together, our results are consistent with the notion that ATP-hydrolysis-driven translocation readily induces CARDs-CARDs interactions between MDA5 proteins, leading to the recruitment of multiple motors in close proximity onto dsRNA.

We further included LGP2 in the MDA5 foci formation experiments. Intriguingly, upon the addition of LGP2, the frequency of MDA5 foci slightly increased by ~ 2-fold while these foci predominantly became immobile (Figures 4C and 4D). In contrast, when LGP2 was substituted with LGP2(E116A), most of the MDA5 foci remained in the translocation state (Figures 4C and 4D). The immobile MDA5 foci displayed significantly greater stability on dsRNA compared with their mobile counterparts, likely because the stationary MDA5-LGP2 oligomers were unable to disassemble from an RNA end (Figures 4E and 4F). These findings highlight the regulatory role of LGP2 in controlling the 1D motions of MDA5 foci, which is consistent with the single-molecule analysis of MDA5-LGP2 interactions (Figure 3).

### Functional assembly of CARDs requires LGP2

While the formation of MDA5 foci certainly relied on CARDs-CARDs interactions, it remained unknown whether the CARDs within MDA5 foci had assembled into tetramers capable of interacting with the adaptor MAVS^11^. We purified and labeled MAVS-CARD with Alexa Fluor 647 (AF647) (Figures S1C and S1D), and co-injected Cy3-MDA5 with AF647-MAVS-CARD proteins to monitor their real-time interactions, probing the formation of MDA5 CARDs tetramers (Figure 5A). In the absence of LGP2, a great number of MDA5 foci were observed translocating along dsRNA, while no MAVS-CARD signal was detected (Figures 5B, 5C and S5A, −LGP2). Addition of LGP2 resulted in numerous MAVS-CARD molecules that were co-localized with immobile MDA5 foci (Figures 5B, 5C and S5A, + LGP2), whereas substitution of LGP2 by LGP2(E116A) significantly decreased MAVS-CARD binding on dsRNA [Figures 5B, 5C and S5A, + LGP2(E116A)]. Further validation revealed that the recruitment of MAVS-CARD required MDA5’s CARDs, as replacing MDA5 with MDA5ΔN resulted in no observable MAVS-CARD binding on dsRNA, even in the presence of LGP2 (Figures 5B, 5C and S5A). These results suggest that the CARDs from translocating MDA5 molecules are unlikely to form tetramers, whereas the MDA5-LGP2 interactions induce the higher-order assemblies of CARDs (Figure 5D).

**Figure 5.**
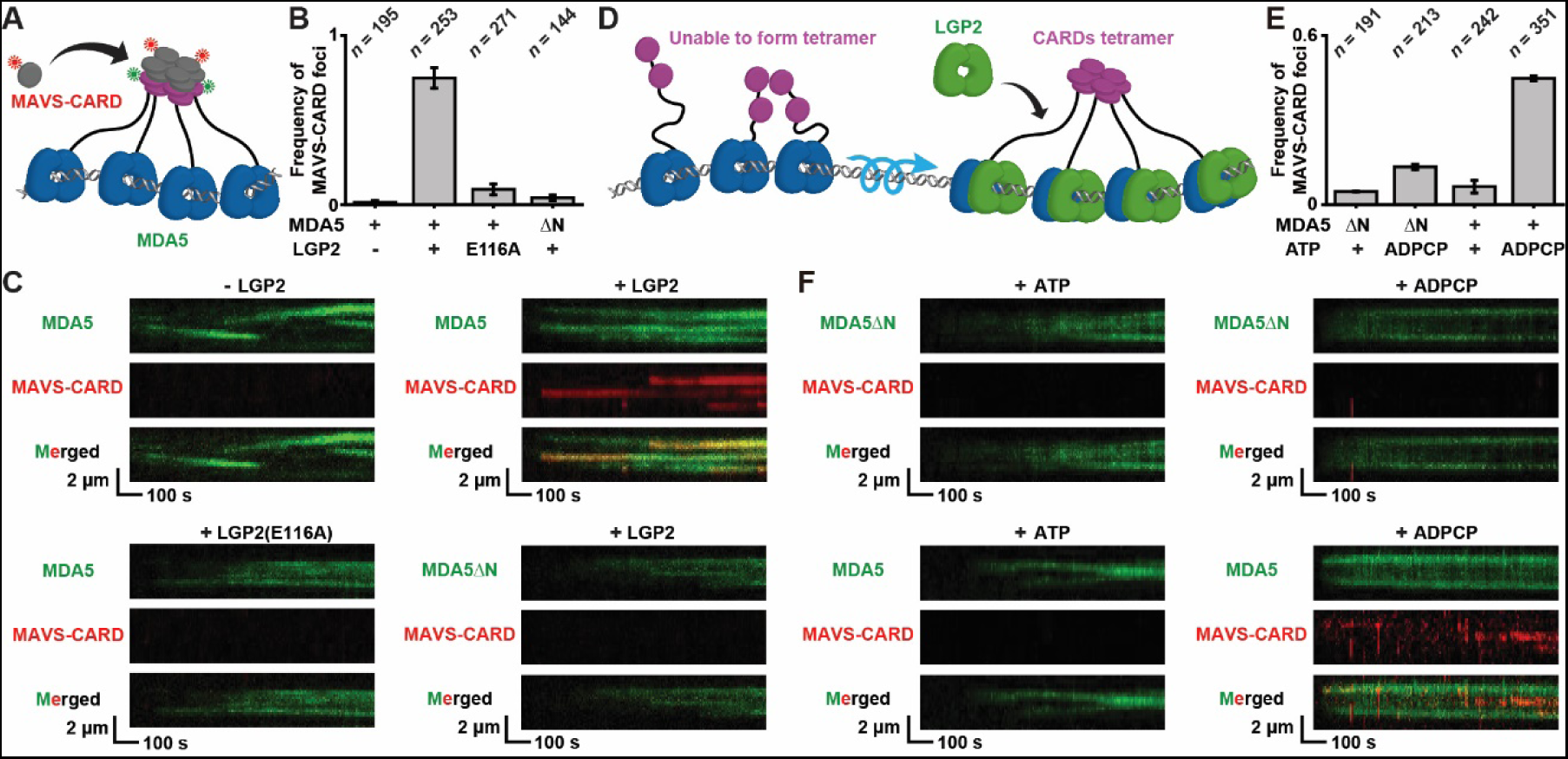
Functional assembly of CARDs requires LGP2. (A) A schematic illustration of the MAVS-CARD recruitment experiment. (B) The frequency of MAVS-CARD foci recruited by MDA5 under various conditions (mean ± s.d.; *n* = number of dsRNA molecules). (C) Representative kymographs showing the recruitments of AF647-MAVS-CARD (100 nM) by Cy3-MDA5 or Cy3-MDA5ΔN (20 nM) under various conditions. The absence or presence of LGP2/LGP2(E116A) (20 nM) is indicated above each set of kymographs. (D) A schematic illustration showing the CARDs from mobile MDA5 molecules are unable to form tetramers, whereas the CARDs from immobile MDA5-LGP2 complexes can assemble into functional tetramers. (E) The frequency of MAVS-CARD foci recruited by MDA5 or MDA5ΔN under various conditions (mean ± s.d.; *n* = number of dsRNA molecules). (F) Representative kymographs showing the recruitments of AF647-MAVS-CARD (100 nM) by Cy3-MDA5 or Cy3-MDA5ΔN (100 nM) under various conditions. The presence of ATP or ADPCP is indicated above each set of kymographs.

The mobile MDA5 foci facilitating CARDs-CARDs interactions (Figures 4A and 4B) while simultaneously suppressing the formation of CARDs tetramers (Figures 5B and 5C) was entirely unexpected. Given that LGP2 appears to interact specifically with the MDA5 ring-like clamp (Figure 3), we envision that this “switch” triggers the functional CARDs assembly by inhibiting the translocation activity of MDA5. To test this hypothesis, MDA5 or MDA5ΔN, along with MAVS-CARD, were co-injected with ATP or ADPCP, followed by single-molecule imaging. Under conditions allowing ATP hydrolysis, both MDA5 and MDA5ΔN motors were found translocating along dsRNA, with no MAVS-CARD foci detected (Figures 5E, 5F and S5B, + ATP). Replacement of ATP with ADPCP completely abolished protein translocation, resulting in the dsRNA being coated by numerous immobile MDA5 or MDA5ΔN molecules (Figures 5E, 5F and S5B, + ADPCP). Interestingly, ADPCP largely enhanced the binding of MAVS-CARD to MDA5 but not to MDA5ΔN, indicating the assembly of CARDs tetramers when MDA5 translocation was impeded (Figures 5E, 5F and S5B, + ADPCP). These findings together suggest it is the translocation of MDA5 that prevents CARDs from forming tetramers, and a fundamental role of LGP2 is to stop MDA5 translocation on dsRNA.

The force generated by ATP hydrolysis seemed to empower the motor to disrupt the biotin-neutravidin binding^45^ (Figure S4C), one of the strongest non-covalent interactions in nature. These observations imply that cellular MDA5 proteins are likely to remain mobile on viral RNAs, with LGP2 being essential for turning the motors off. To validate LGP2’s role in cellular MDA5 signaling, we measured interferon mRNA levels in HEK293T (293T) cells following stimulation with high-molecular-weight (HMW) poly(I:C), a dsRNA mimetic. Due to the poor expression of endogenous RLR proteins, MDA5 signaling activity was relatively weak in wild-type 293T cells^22^ (Figures 6A and 6B). Overexpression of MDA5 slightly enhanced poly(I:C)-induced interferon production, while LGP2 supplementation showed no observable effect (Figures 6A and 6B). In contrast, co-expression of MDA5 and LGP2 largely elevated interferon mRNA levels, indicating LGP2’s potentials to activate MDA5 signaling in cells. To further confirm the LGP2’s essential roles in MDA5-mediated antiviral immune response, we generated MDA5 or LGP2 knockout A549 cells using CRISPR/Cas9 (referred to as ΔMDA5 and ΔLGP2, respectively, Figure 6C). Interferon mRNAs were subsequently monitored following encephalomyocarditis virus (EMCV) infections. Compared with wild-type A549, both ΔMDA5 and ΔLGP2 cells exhibited significant deficiencies in interferon productions (Figure 6D). These results align with earlier *in vivo* and cellular studies that have demonstrated an indispensable role of LGP2 in MDA5-dependent immune response to both virus infections and endogenous RNAs^4,46,47^.

**Figure 6.**
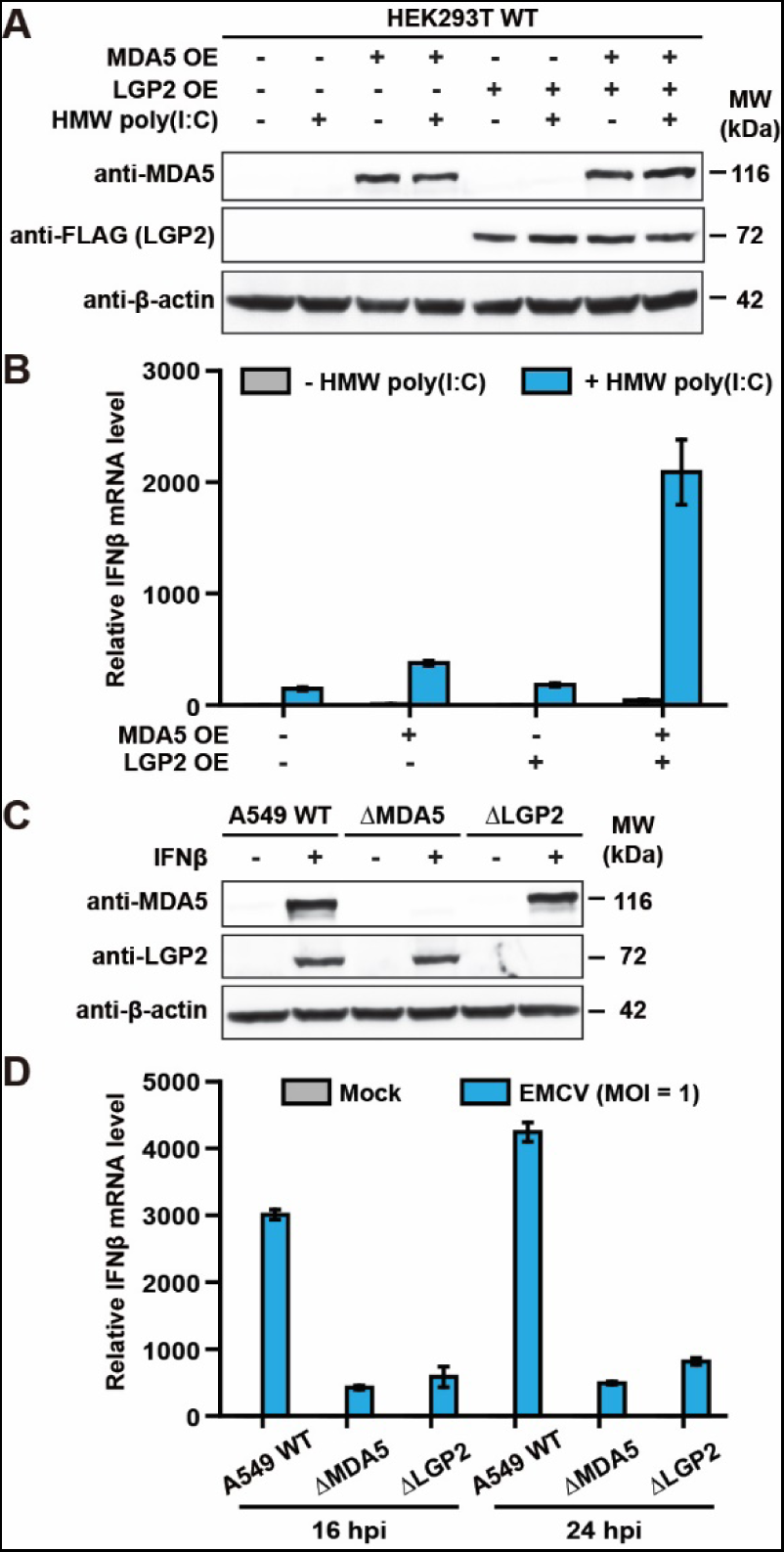
LGP2 is indispensable for MDA5-dependent immune response. (A) Immunoblotting showing the overexpression (OE) of MDA5/LGP2 proteins in 293T cells. Proteins were detected using indicated antibodies. For poly(I:C) stimulation, cells were transfected with 1 μg/mL HMW poly(I:C) for 24 h. (B) Relative IFNβ mRNA levels showing the MDA5 signaling activities in wild-type and MDA5/LGP2-overexpressed 293T cells. IFNβ productions were induced upon HMW poly(I:C) stimulation. (C) Immunoblotting showing the successful knockouts of MDA5/LGP2 in A549 cells. Proteins were detected using indicated antibodies. Endogenous MDA5/LGP2 expression was induced by interferon stimulation. (D) Relative IFNβ mRNA levels showing the MDA5 signaling activities in wild-type A549, ΔMDA5 and ΔLGP2 cells. IFNβ productions were induced upon EMCV infections. MOI: multiplicity of infection, hpi: hours post-infection.

## DISCUSSION

It has been known for decades that MDA5 is capable of measuring the length of a dsRNA molecule by ATP hydrolysis^1,48,49^. Mutations that interfere with MDA5 ATPase activity cause autoimmune diseases due to impaired ability to distinguish between self and non-self RNAs^50–52^. Previous biochemical assays have shown that ATP hydrolysis triggered the release of MDA5 molecules from dsRNA^25^, while these results were initially assumed to support an ATP-regulated assembly and/or disassembly model for MDA5 filament^23^. Here our single-molecule results undoubtedly demonstrate that the role of ATP hydrolysis is to drive the protein translocation. The dissociation of MDA5 motors from a substrate is likely attributed to the discontinuous backbone at the dsRNA end (Figures 1C, 4A and S4C). These observations imply that MDA5 can undergo long-distance translocation along viral dsRNA, enabling the motor to associate with LGP2 for signaling (Figure S6, top). In contrast, the translocation of MDA5 on self RNA may be short-lived and insufficient for the recruitment of downstream components (Figure S6, bottom). Therefore, a fundamental role of rotation-coupled 1D translocation is to interrogate the integrity and measure the length of a dsRNA substrate. When ATP hydrolysis is inhibited, MDA5 appears to bind immobile to internal sites of a dsRNA (Figures 1C and 5F), leading to the formation of abnormal MDA5 oligomers that might activate downstream signaling on self RNA.

Probably the most surprising finding in our study is the incapability of mobile MDA5 foci to form functional oligomers. The detailed mechanism behind this observation remains elusive due to the limited spatial resolution of smTIRF. One might speculate that an MDA5 molecule also maintains an auto-repressed state, wherein LGP2 binding rearranges MDA5 conformation to expose the CARDs. However, the observations of CARDs-CARDs interactions without LGP2 (Figures 4A and 4B), along with the induction of tetramer formation by ADPCP (Figures 5E and 5F), largely dismiss this idea. One possible explanation is that the MDA5 motor moves in a rotation-coupled manner, creating a scaffold for the flexible IDR linker between CARDs and helicase domain to wrap around the dsRNA in concert with natural helical motions (Figure S6, top). As a result, this configuration significantly constrain the motility of the CARDs, thereby diminishing their ability to come into close proximity and form tetramers (Figure S6, top). It appears that the 1D translocation can completely suppress the spontaneous CARDs assembly, even under high protein concentrations where substantial amounts of MDA5 motors have been loaded onto a single dsRNA (Figures 5E and 5F, + ATP). Notably, although the MAVS-CARD foci were utilized to probe the formation of MDA5 CARDs tetramers in our single-molecule study, it does not necessarily imply that cellular MAVS-MDA5 interactions occur on dsRNA. Both poly-ubiquitin and E3 ligase TRIM65 are likely to assist in stabilizing CARDs oligomers in cells, and MDA5 may be released from dsRNA as functional tetramers and delivered to MAVS^11,22,53,54^.

While the existing model proposes that multiple MDA5 proteins assemble into a filament relying on helicase-helicase stacking between adjacent molecules^23^, no filament or filament-like oligomers of MDA5(Q57E) or MDA5ΔN were observed at the protein concentrations used in smTIRF (3 – 100 nM, Figures 4A, 4B, 5C and 5F). In contrast, direct interactions between MDA5ΔN and LGP2 were detected at sub-nanomolar protein concentrations (0.5 nM, Figures 3E and S4B). These results illustrate that the MDA5ΔN-LGP2 interactions are considerably stronger than the helicase-helicase contacts between two MDA5ΔN molecules, a finding also supported by a recent molecular modelling study^45^. At an elevated concentration (15 nM), wild-type MDA5 indeed formed foci that moved along dsRNA, showing clear dependence on CARDs rather than helicase-helicase interactions (Figures 4A and 4B). Whether LGP2 binding induced filament formation in an MDA5 foci remains unclear (Figure 4C); however, all the MDA5 motors appeared to be turned off prior to oligomerization. It is possible that the formation of MDA5 filaments requires higher protein concentrations, typically occurring during the late phase of antiviral response, as interferon induces a substantial increase in MDA5 protein expression^55,56^. On the contrary, cells must be capable of responding to viral infections before MDA5 accumulation. Thus, the initiation of antiviral immune response is LGP2-dependent^4,46,47^ (Figures 6B and 6D), likely owing to the supreme signaling activities of MDA5-LGP2 complexes (Figure 5).

The helicase domains and CTDs of RLR sensors exhibit similar sequence conservation^57^, yet they may have distinct biological functions. We found that MDA5 recognized dsRNA backbone internally and initiated processive translocation, whereas long-distance movement of RIG-I had not been observed (Figure S3). It is likely that RIG-I motors translocated along dsRNA with low processivity, events of which might not be detectable under the spatial resolution of our imaging system (200 – 400 nm)^32^. While it is clear that the translocation of MDA5 on long dsRNA recruits LGP2 and activates downstream signaling (Figure 5), the physiological roles of RIG-I translocation continue to be uncertain and require further investigations. LGP2, although it shares a domain arrangement similar to that of MDA5 and RIG-I^58,59^, has been shown here to lack stable binding to dsRNA in the presence of ATP (Figure 2). We cannot exclude the possibility of LGP2 performing short-distance translocation on dsRNA^60^, however, our results strongly suggest that the helicase functions as a mediator to connect RNA recognition with CARDs assembly, at least in the MDA5 signaling pathway.

Collectively, our findings reveal that the assembly of oligomers may be greatly influenced by protein motions on nucleic acids, emphasizing the orchestrated nature of RNA-protein interactions that employ 1D movement to complete the progressions of innate immune response.

## ACKNOWLEDGMENTS

We would like to thank Dr. Richard Fishel (The Ohio State University) and Dr. Dong Gao (CAS Center for Excellence in Molecular Cell Science) for many helpful insights and discussions, and Dr. Zhuo Yang (Chemical Biology Core Facility of CAS Center for Excellence in Molecular Cell Science) for technical support. This work was supported by the Chinese Academy of Sciences (CAS) (YSBR-009), the Strategic Priority Research Program of the Chinese Academy of Science (XDB0570000), the National Key R&D Program of China (2021YFA1300503), and the National Natural Science Foundation of China (32071283) to J. Liu.

## AUTHOR CONTRIBUTIONS

X.-P.H., M.R. and J.L. designed the experiments; X.-P.H. and S.-R.W. purified and labeled the proteins; X.-P.H. and S.-R.W. performed the single molecule studies; M.R. performed the cellular MDA5 signaling assays; X.-P.H., M.R., S.-R.W. and J.L. analyzed the data; S.-Q.Z. performed the molecular modeling; X.-P.H., M.R., F.-J.H., L.-L.C. and J.L. wrote the paper and all authors participated in critical discussions.

## METHODS

### Cell culture

HEK293T (ATCC) cells were cultured in Dulbecco’s modified Eagle’s medium (DMEM; Gibco) supplemented with 10% fetal bovine serum (Gibco) and 100 U/mL of penicillin/streptomycin (Gibco). A549 (ATCC) cells were cultured in Ham’s F-12K (Kaighn’s) medium (Gibco) supplemented with 10% fetal bovine serum (Gibco) and 100 U/mL of penicillin/streptomycin (Gibco). Both HEK293T and A549 cells were grown at 37 °C in a 5% CO_2_ humidified atmosphere. SF9 insect cells were cultured in Sf-900™ II SFM medium (Gibco) supplemented with 1 x GlutaMax (Gibco) and 10 μg/mL Gentamicin (ABCONE), and were grown in suspension cultures at 27 °C while shaking at 140 rpm.

### Plasmid construction, recombinant protein labeling and purification

The human MDA5, MDA5(Q57E), MDA5ΔN (residues 287-1025), LGP2, LGP2(K30A), LGP2(E116A) and RIG-I proteins were labeled using sortase-mediated peptide ligation^61,62^. The RLR genes were amplified by PCR (Table S1), digested with BamHI and KpnI (for MDA5 and LGP2), or NotI and XhoI (for RIG-I), and inserted into pFastbac1 baculovirus expression plasmid. Hexa-histidine (his_6_) and sortase recognition sequence (srt, LPETG) were introduced onto the the N-terminus of MDA5 and RIG-I proteins, or C-terminus of LGP2 protein. The MDA5(Q57E), LGP2(K30A), and LGP2(E116A) mutations were generated using the KOD-Plus-Mutagenesis Kit (Toyobo, cat. no. SMK-101). Two serine residues separated the his_6_ and srt, and these tags were separated from the RLR proteins by two or four glycine residues. MDA5 and LGP2 knockout A549 cells were generated by CRISPR/Cas9. Single guide RNAs (sgRNAs, Table S1) for MDA5 or LGP2 were cloned into a CRISPR/Cas9-based vector with a puromycin resistance selection marker. To overexpress MDA5 and LGP2 in 293T cells, human MDA5 and LGP2 genes were inserted into pCMV plasmids. A Myc tag was introduced into the N-terminus of MDA5, and a 3 × Flag tag was introduced into the C-terminus of LGP2. All the plasmid constructs were amplified in *E. coli* DH5α and verified by DNA sequencing.

The MDA5, LGP2 and RIG-I proteins were overexpressed in SF9 insect cells using the Bac-to-Bac Baculovirus Expression System (Thermo Fisher Scientific). All purification procedures were carried out at 4 °C. SF9 cells were harvested and suspended in Lysis Buffer (25 mM Tris-HCl pH 7.5, 500 mM NaCl, 20% glycerol, and 20 mM imidazole). Cells were thawed on ice, passed through a 25 G needle five times, and centrifuged at 50,000 x g for 1h. The supernatant was loaded onto a Ni-NTA (Qiagen) column, washed with Buffer A (25 mM Tris-HCl pH 7.5, 500 mM NaCl, 20 % glycerol and 20 mM imidazole), and eluted with Buffer B (25 mM Tris-HCl pH 7.5, 500 mM NaCl, 20 % glycerol and 200 mM imidazole). Peak fractions containing recombinant proteins were pooled and dialyzed against Labeling Buffer (50 mM Tris-HCl pH 7.5, 150 mM or 500 mM NaCl and 20 % glycerol). To label MDA5 and RIG-I, recombinant proteins were incubated with an improved version of sortase^63^ and peptides (Cy3/Cy5-CLPETGG, purchased from ChinaPeptides Co.,LTD) as described^40^ (protein: sortase: peptide in the molar ratio of 1:2:5). To label LGP2, recombinant proteins were incubated with sortase and peptides (GGGC-Cy3/Cy5, purchased from ChinaPeptides Co.,LTD). The reaction was incubated for 1 h on ice and loaded onto a Ni-NTA column. Flow through proteins were collected and diluted with Buffer C (25 mM Tris-HCl pH 7.5, 1 mM DTT, 20 % glycerol and 0.1 mM EDTA) to reduce NaCl concentration within the buffer (~ 100 mM). The proteins were then loaded onto a heparin column, washed extensively with Buffer C plus 100 mM NaCl and eluted with Buffer C plus 1 M NaCl. For MDA5 and RIG-I, further contaminants were separated by size-exclusion chromatography (Superdex 200 10/300 GL, GE Healthcare). Protein-containing fractions were dialyzed against Storage Buffer (25 mM Tris-HCl pH 7.5, 150 mM or 500 mM NaCl, 1 mM DTT, 0.1 mM EDTA, and 20 % glycerol) and frozen at −80 °C.

A MAVS-CARD protein containing three mutations (D23K, E26K, and E80K) was used for single-molecule imaging. These mutations have been reported to increase protein solubility without disrupting MDA5-MAVS interactions^11,64^. A pET47b plasmid encoding MAVS-CARD (residues 1-97)^65^ with an N-terminal His-tag and a C-terminal SNAP-tag was a gift from Sun Hur lab. The MAVS-CARD(D23K/E26K/E80K) (referred as MAVS-CARD in the following text and main text) were generated using the KOD-Plus-Mutagenesis Kit. After transformation with the MAVS-CARD expression plasmid, a single colony of BL21(DE3) cell was diluted into 1 L of LB containing 50 μg/ml kanamycin. At OD_600_ = 0.4, the growth temperature was decreased to 18 °C and the expression protein was induced by the addition of IPTG (0.2 mM) at 18 °C for 16 h. Cells were collected and resuspended in Freezing Buffer (25 mM Tris-HCl pH 7.5, 500 mM NaCl, 20 % glycerol and 20 mM imidazole). Cell pellets were freeze-thawed three times, sonicated twice, and centrifuged at 50,000 × g for 1 h. The supernatants were then loaded onto a Ni-NTA column, washed with Buffer A, Buffer D (25 mM Tris-HCl pH 7.5, 500 mM NaCl, 20 % glycerol and 60 mM imidazole), and eluted with Buffer B. Fractions containing MAVS-CARD were pooled, and incubated with SNAP-Surface Alexa Fluor 647 substrate (New England Biolabs) in 1 mM DTT at 4 °C for overnight. After labeling, the protein was diluted with Buffer E (25 mM Tris-HCl pH 7.5, 500 mM NaCl, 20% glycerol) to reduce imidazole concentration within the buffer (~ 20 mM). Proteins were then loaded onto the Ni-NTA column, washed with Buffer A and eluted with Buffer B. Protein-containing fractions were dialyzed against Storage Buffer and frozen at −80 °C. Given the observations that purified MAVS-CARD proteins exist in short filamentous structures, a denaturing-refolding approach was employed to obtain monomeric MAVS-CARD as described^65^. In brief, MAVS-CARD was denatured in 6 M guanidinium hydrochloride (GuHCl) at 37 °C for 30 min, followed by dialysis in Refolding Buffer (20 mM Tris-HCl, pH 7.5, 500 mM NaCl, 0.5 mM EDTA and 2 mM DTT) at 4 °C for 1 h, and was immediately used for single-molecule imaging.

The concentrations and labeling efficiencies of all recombinant proteins were determined by measuring protein absorbance at 280 nm and Cy3/Cy5 absorbance at 550/650 nm, respectively (Table S2).

### ATPase analysis

The ATPase activities of MDA5 and LGP2 were measured utilizing an ATPase/GTPase Activity Assay Kit (Sigma). The analysis was carried out with 200 nM protein and 100 ng RNA substrate (for ssRNA: ssRNA31, Table S1; for dsRNA: 11.6-kb dsRNA) in a 40 μL reaction mixture comprised of 20 mM Tris-HCl (pH 7.5), 150 mM NaCl, 1 mM ATP, 5 mM MgCl_2_, and 1 mM DTT. For MDA5, the reactions were performed at 23 °C for 0, 1, 2, 3, and 4 min followed by quenching with 200 μL of malachite green reagents. For LGP2, the reactions were performed at 23 °C for 0, 10, 20, 30, and 40 min followed by quenching with 200 μL of malachite green reagents. Samples were incubated at 23 °C for additional 30 min. Free phosphate was determined by measuring absorbance at 620 nm using a microplate reader Eon (Bio Tek). Data were fit to a linear function to calculate the rates of ATP hydrolysis and the turnover numbers (k_cat_).

### Preparation of 11.6-kb dsRNA

Two complementary 11.6-nt ssRNA strands were produced by SP6 in vitro transcription, followed by annealing together along with two biotin-labeled ssRNA linkers to generate an 11.6-kb dsRNA substrate (Figure S1A). In brief, two plasmids containing the SP6 promoter and sense/anti-sense strand sequences were constructed, amplified in E. coli DH5α and verified by DNA sequencing. DNA templates were then linearized by KpnI (for sense strand) or BamHI (for anti-sense strand) digestion, and purified by phenol-chloroform extraction and ethanol precipitation. RNAs were transcribed in vitro using SP6 RNA Polymerase (Thermo Fisher Scientific), followed by phenol-chloroform extraction and ethanol precipitation. Sense and anti-sense RNA strands (100 nM for each), along with RNA Linker 1 and RNA Linker 2 (10 µM for each; Table S1) were mixed in Annealing Buffer (20 mM Tris-HCl pH 7.5, 50 mM NaCl, 1 mM EDTA), denatured at 95 °C for 10 min, then cooled down slowly to 23 °C. The resulting product was separated on a 0.75 % agarose gel and the 11.6-kb dsRNA was excised and purified using Agarose Gel DNA Extraction Kit (Takara).

The 11.6-kb dsRNA substrate for RIG-I binding experiments was prepared using a similar protocol, with the exception that the binding site for RNA Linker 1 on the sense strand was removed. As a result, the dsRNA was tethered at one end only via biotin-neutravidin, with a 5’-ppp located at the other end (Figure S3A).

### Single-molecule imaging buffers and experiment conditions

The single-molecule Imaging Buffer contains 20 mM Tris-HCl (pH 7.5), 2 mM DTT, 0.2 mg/mL acetylated BSA (Molecular Cloning Laboratories), 0.0025% P-20 surfactant (GE healthcare), 2 mM ATP (unless stated otherwise), 1.5 mM MgCl_2_ and 100 mM NaCl. All single-molecule experiments were carried out at 23 °C.

### Single molecule total internal reflection fluorescence (smTIRF) microscopy

All the single molecule total internal reflection fluorescence (smTIRF) data were acquired on a custom-built prism-type TIRF microscope established on the Olympus microscope body IX73^40^. Fluorophores were excited using the 532 nm for green and 637 nm for red laser lines built into the smTIRF system. Image acquisition was performed using an EMCCD camera (iXon Ultra 897, Andor) after splitting emissions by an optical setup (OptoSplit II emission image splitter, Cairn Research). Micro-Manager image capture software was used to control the opening and closing of a shutter, which in turn controlled the laser excitation.

The 11.6-kb dsRNA (50 pM) in 500 μL T50 buffer (20 mM Tris-HCl, pH 7.5, 50 mM NaCl) was injected into a custom-made flow cell chamber and stretched by laminar flow (300 μL/min). The stretched dsRNA was anchored at both ends onto a neutravidin-coated, PEG passivated quartz slide surface, and the unbound dsRNA was flushed by similar laminar flow. Unless stated otherwise, a 5000-ms frame rate with 300-ms laser exposure time was used to minimize photo-bleaching during single-molecule imaging. The dsRNA was located by staining with SYBR Gold (0.2 X, Invitrogen) after real-time recording.

To examine MDA5 translocation on dsRNA, 3 nM Cy3-MDA5 in Imaging Buffer was introduced into the flow cell chamber. To examine LGP2 binding on dsRNA, 60 nM Cy3-LGP2 with or without 60 nM unlabeled MDA5 in Imaging Buffer was introduced into the flow cell chamber. The interactions between MDA5/LGP2 proteins and dsRNA were monitored in real-time for 25 min in the absence of flow. To examine the MDA5-LGP2 interactions on dsRNA, 3 nM Cy3-MDA5 and 3 nM Cy5-LGP2, or 0.5 nM Cy3-MDA5ΔN and 0.5 nM Cy5-LGP2 proteins in Imaging Buffer were introduced into the flow cell chamber and protein-protein interactions monitored in real-time for 25 min in the absence of flow.

To examine RIG-I binding on dsRNA, a single-biotin-labeled 11.6-kb dsRNA was injected into the flow cell chamber. After 20 min incubation, 5 nM Cy5-RIG-I protein in Imaging Buffer was introduced. The interactions between RIG-I and dsRNA were monitored in real-time with a flow rate at 100 μL/min. A 300-ms frame rate with 300-ms laser exposure time was used to image RIG-I on dsRNA.

To examine the formation of MDA5 foci, 15 nM Cy3-MDA5 in Imaging Buffer was introduced into the flow cell chamber. To examine the formation of MDA5 foci in the presence of LGP2, 10 nM Cy3-MDA5 with or without 10 nM unlabeled LGP2 proteins in Imaging Buffer were introduced into the flow cell chamber. The MDA5 molecules on dsRNA were then monitored in real-time for 25 min in the absence of flow. To measure the survival probability of MDA5 foci, 10 nM Cy3-MDA5 with or without 10 nM unlabeled LGP2 proteins in Imaging Buffer were first introduced into the flow cell chamber. After 17 min incubation, the flow cell was flushed with Imaging Buffer and the MDA5 foci on dsRNA were monitored in real-time for 21 min in the absence of flow.

To examine the interactions between MDA5 and MAVS-CARD, Cy3-MDA5 (20 nM), unlabeled LGP2 (20 nM) and AF647-MAVS-CARD (100 nM) in Imaging Buffer were introduced into the flow cell chamber. To examine the interactions between MDA5 and MAVS-CARD in the presence of ADPCP, Cy3-MDA5 (100 nM) and AF647-MAVS-CARD (100 nM) in Imaging Buffer were introduced into the flow cell chamber and protein-protein interactions were monitored in real-time for 17 min in the absence of flow.

### Position determination on dsRNA

To determine the positions of MDA5 and RIG-I proteins on RNA, the 11.6-kb dsRNA was stained with SYBR Gold (0.2 X, Invitrogen). The left (*P_L_*) and the right (*P_R_*) end positions of the dsRNA were determined by plotting the fluorescent intensities along the length of the stained RNA as previously described ^33^. Horizontal positions of protein particles (*P_P_*) on dsRNA were tracked by DiaTrack 3.05 (Sydney, Australia), where the particle intensities were fit to a two-dimensional Gaussian function to obtain their positions with sub-pixel resolution. The positions were then converted to lengths in bp by the following equation: 11,645 bp × (*P_P_* − *P_L_*) / (*P_R_* − *P_L_*), in which 11,645 bp is the length of the dsRNA. A 1000 bp (~1 pixels) binning size was used to construct the position histograms.

### Data analysis of TIRF imaging

All kymographs were generated along the dsRNA by a kymograph plugin in ImageJ (J. Rietdorf and A. Seitz, EMBL Heidelberg). For studies involving Cy3-MDA5 and Cy5-LGP2, or Cy3-MDA5 and AF647-MAVS-CARD, fluorescent molecules in two channels were co-localized using a custom written MATLAB script. Particles were tracked using DiaTrack 3.05 to obtain single-molecule fluorescent intensities and trajectories.

To determine the frequency of MDA5 translocation on dsRNA, single-molecule movies were recorded for 25 min and the translocating MDA5 molecules with a minimum lifetime of 40 s were counted as the number of MDA5 translocation (*N*_MDA5-trans_). To determine the frequency of LGP2-coated dsRNA, single-molecule movies were recorded for 25 min and the string-like fluorescent LGP2 molecules with a minimum length of 7 pixels were counted as the number of LGP2-coated dsRNA (*N*_LGP2-dsRNA_).

To determine the frequency of MDA5ΔN translocation and stop events in the presence/absence of LGP2, single-molecule movies were recorded for 25 min and the translocating MDA5ΔN molecules with a minimum lifetime of 40 s were counted as the number of MDA5ΔN translocation events (*N*_MDA5ΔN-trans_). The MDA5ΔN molecules undergoing 1D translocation and subsequently transitioning into immobile state were counted as the number of MDA5ΔN stop events (*N*_MDA5ΔN-stop_).

To determine the frequency of mobile and immobile MDA5 foci in the presence/absence of LGP2, single-molecule movies were recorded for 25 min. The MDA5 molecules with increased fluorescent intensities (> 10-fold of the initial intensity of a single MDA5) during 1D translocation were counted as the number of mobile MDA5 foci (*N*_MDA5-mobile_). The immobile MDA5 molecules with increased fluorescent intensities (> 10-fold of the initial intensity of a single MDA5) were counted as the number of immobile MDA5 foci (*N*_MDA5-immobile_). To determine the frequency of MAVS-CARD foci, single-molecule movies were recorded for 17 min and the MAVS-CARD molecules with a minimum lifetime of 50 s were counted as the number of MAVS-CARD foci (*N*_MAVS-CARD_).

Following the real-time single-molecule recording, the number of dsRNA molecules (*N*_RNA_) was determined by SYBR Gold staining. The frequencies of MDA5 translocation (*F*_MDA5-trans_), LGP2-coated dsRNA (*F*_LGP2-dsRNA_), MDA5ΔN translocation event (*F*_MDA5ΔN-trans_), MDA5ΔN stop event (*F*_MDA5ΔN-stop_), mobile MDA5 foci (*F*_MDA5-mobile_), immobile MDA5 foci (*F*_MDA5-immobile_) and MAVS-CARD foci (*F*_MAVS-CARD_) were calculated using the following equations that also included corrections for labeling efficiencies of the proteins (the numbers in the denominator, Table S2):

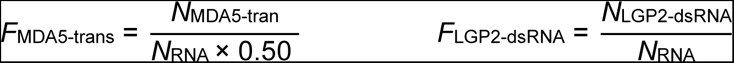

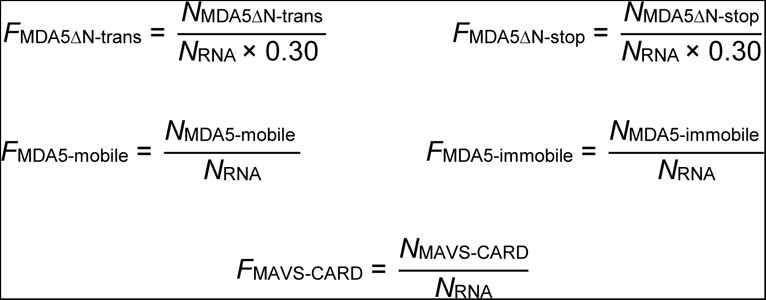

All single molecule frequency studies were performed at least two separate times.

### Survival probability analysis

To plot the survival probability of MDA5ΔN molecules on dsRNA, all events were synchronized in time, with the initial event count set to 1. MDA5ΔN dissociation was quantified in 240-s time bins. To plot the survival probability of MDA5 foci on dsRNA, the number of MDA5 foci at the beginning of each movie was set to 1, and MDA5 dissociation was quantified in 250-s time bins.

### Translocation rates analysis

The association (*P*_A_) and dissociation (*P*_D_) positions of MDA5 translocation were determined by particle trajectories on dsRNA. The time intervals (*T*) between association and dissociation events were extracted from kymograph. Translocation rates (*R*) were calculated using the following equation: 11,645 bp × (*P*_A_ − *P*_D_) / (*T* × 2.5 µm), where 2.5 µm is the average length of dsRNA observed by smTIRF microscopy (Figure 1B). Absolute values were used to construct histograms of translocation rates, except for the MDA5ΔN-LGP2 complexes, due to their rates being too close to zero.

### Binning method

All binned histograms were produced by automatically splitting the data range into bins of equal size by using the Origin program.

### Molecular modeling of an MDA5-LGP2 complex

The model of an MDA5-LGP2 complex was predicted by AlphaFold2 within ColabFold (version 1.5.2)^66^, employing the alphafold2_multimer_v3 model and Amber minimization, and the MDA5-LGP2-dsRNA model was obtained using AlphaFold3 server^67^. E116 on LGP2 was at the interface with the helicase domain of MDA5 in both models.

### MDA5/LGP2 overexpression and poly(I:C) stimulation

MDA5/LGP2-expressing plasmids or HMW poly(I:C) (InvivoGen) transfections were carried out using Lipofectamine 2000 Reagent (Invitrogen) according to the manufacturer’s protocols. For MDA5/LGP2 overexpression in 293T, cells were seeded in a 12-well plate and transfected with 500 ng of the pCMV-Myc-MDA5 and/or pCMV-LGP2-3 × Flag plasmids, respectively. After 24 h, cells were further transfected with 1 μg/mL HMW poly(I:C) and incubated for 24 h.

### Generation of ΔMDA5 and ΔLGP2 cells

To generate MDA5 or LGP2 knockout A549, cells were initially seeded in a 6-well plate and subsequently transfected with 2 μg of either the pC338-MDA5 sgRNA or pC338-LGP2 sgRNA plasmids, respectively. Following incubation for 36 h, the medium was replaced, and 2 μg/mL of puromycin was introduced for selection. After additional 48 h, single cells were seeded into a 96-well plate and expanded for screening through sequencing and immunoblotting.

To verify the knockouts of A549 cells, wild-type A549, ΔMDA5 and ΔLGP2 cells were initially seeded in a 6-well plate, the medium was replaced, and 5 ng/mL of IFNβ (Peprotech) was introduced. After 24 h incubation, cells were collected for immunoblotting.

### EMCV infection assay

Wild-type A549, ΔMDA5 and ΔLGP2 cells were seeded in a 48-well plate followed by the addition of EMCV (1 MOI) into the culture medium. Subsequently, total RNAs were collected after 16 or 24 hours for further analysis.

### RT-qPCR

Total RNAs were extracted using RNAiso Plus reagent (Takara) and cDNA was synthesized using PrimeScript™ RT reagent Kit with gDNA Eraser (Perfect Real Time) (Takara) according to the manufacturer’s instructions. Interferon expressions were measured using TB Green® Premix Ex Taq™ (Tli RNaseH Plus) reagent (Takara) on a QuantStudio™ 6 Flex (Applied Biosystems), where β-actin was examined as an internal control for normalization. Primers for qRT-PCR were listed in Table S1.

## SUPPLEMENTAL FIGURE LEGENDS

**Figure S1.**
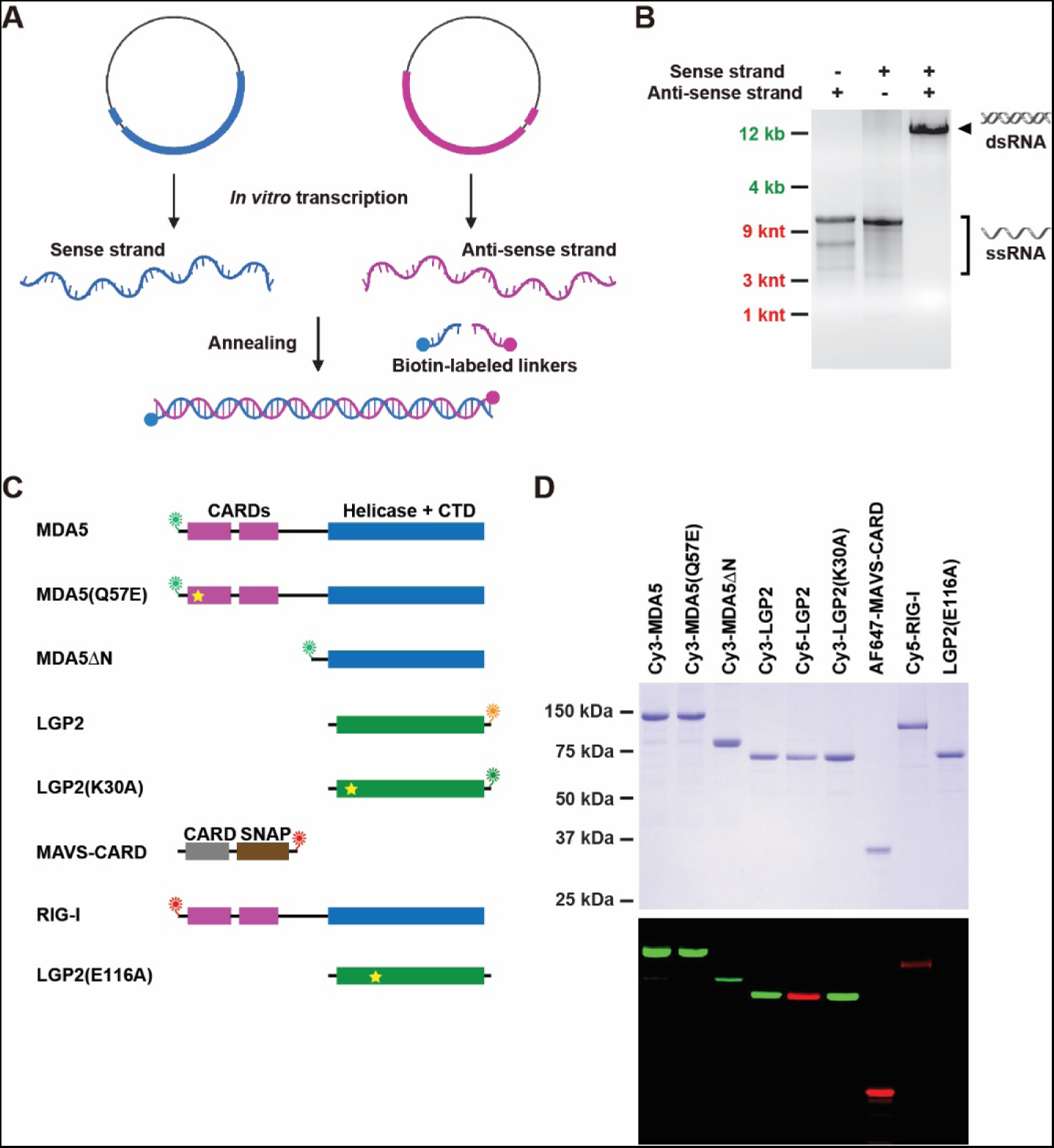
The construction of 11.6-kb dsRNA and fluorophore-labeled proteins used in smTIRF microscopy. (A) A schematic illustration for the construction of an 11.6-kb dsRNA. Sense and anti-sense strands of ssRNA were annealed with two biotin-labeled RNA linkers to generate 11.6-kb dsRNA. (B) Agarose gel (0.75 %) showing the sense strand, anti-sense strand of ssRNA and the annealed 11.6-kb dsRNA. (C) A schematic illustration of labeled and unlabeled proteins used in single-molecule studies. (D) Coomassie stained (top) and fluorescent (bottom) images of SDS-PAGE gels showing the purified proteins.

**Figure S2.**
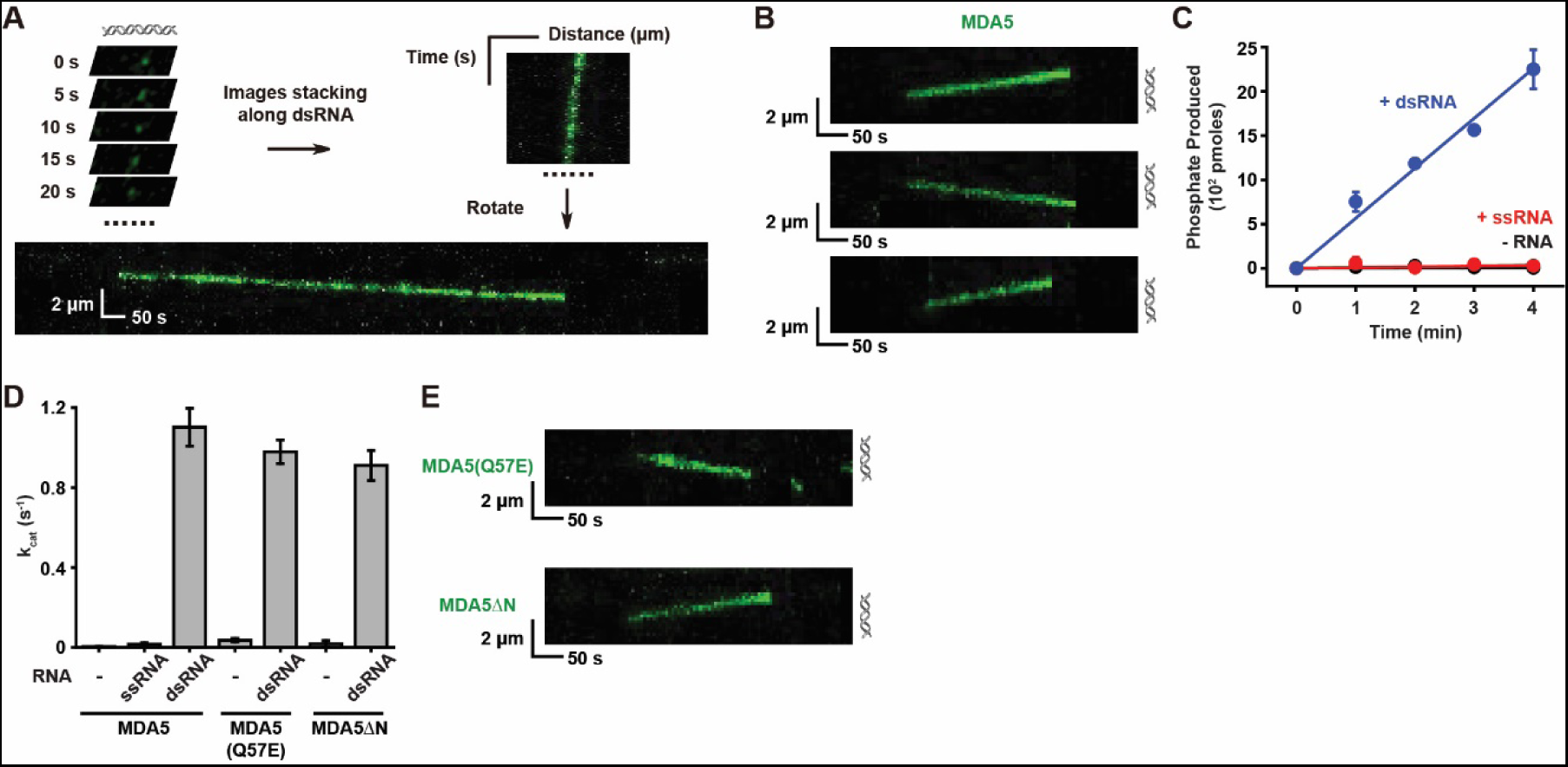
ATPase activities, representative kymographs, translocation rates, and processivity of MDA5 proteins. (A) An illustration of the kymograph construction of a single MDA5 motor on 11.6-kb dsRNA. (B) Representative kymographs showing the translocation of three independent Cy3-MDA5 motors on 11.6-kb dsRNA in the presence of ATP. (C) ATP hydrolysis of MDA5 measured at various time using different RNA substrates. Circles represent individual numbers from at least three independent experiments (error bars: mean ± s.e.). A linear function was fit to the data to derive the turnover number (k_cat_) of MDA5 ATPase (mean ± s.e.). (D) The turnover numbers (k_cat_) of MDA5, MDA5(Q57E) and MDA5ΔN ATPase using different RNA substrates (error bars: mean ± s.e.). (E) Representative kymographs showing the translocation of MDA5(Q57E) and MDA5ΔN on dsRNA.

**Figure S3.**
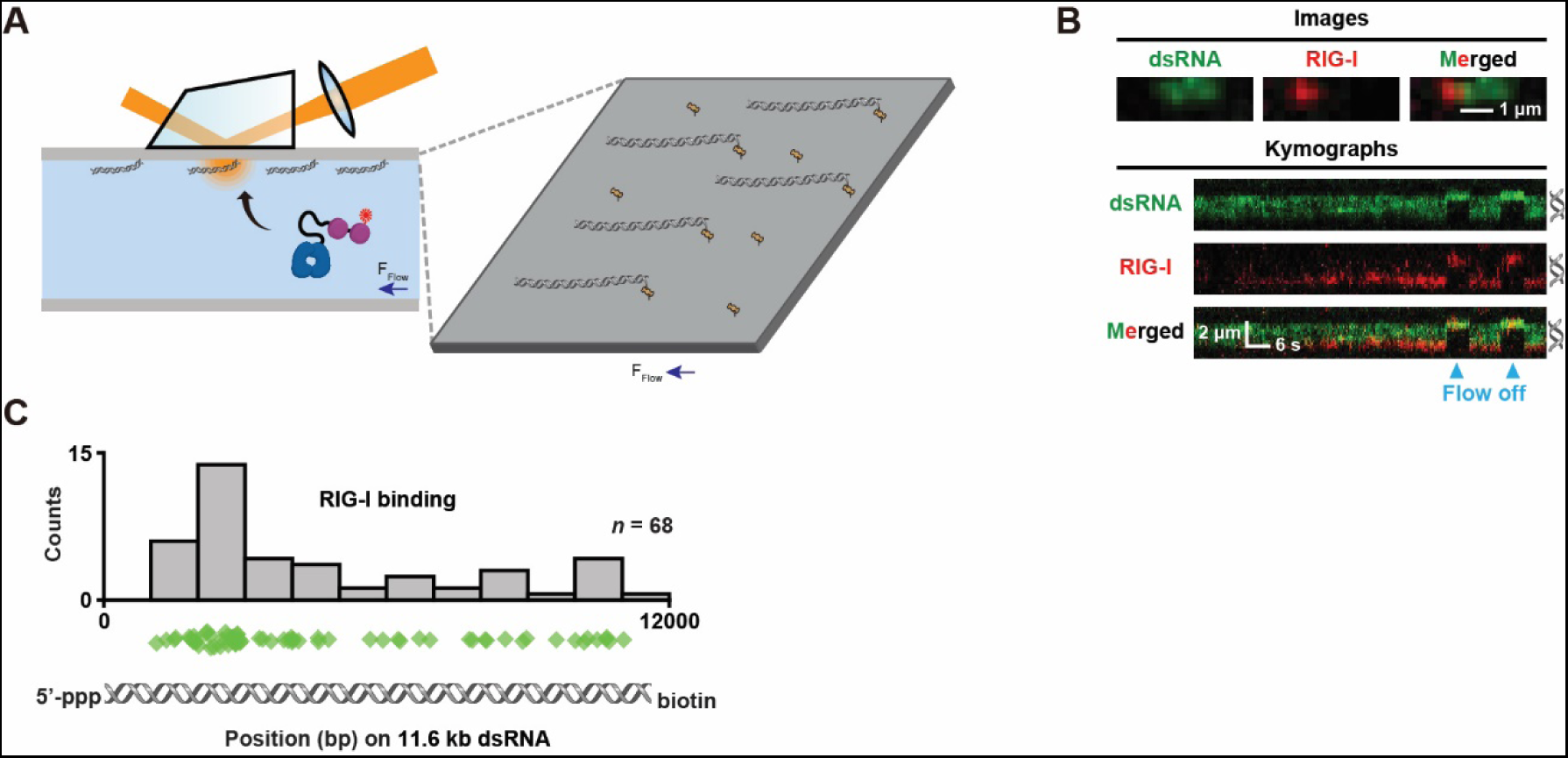
RIG-I recognizes dsRNA end containing 5’-ppp. (A) An illustration of imaging Cy5-RIG-I on a single-end-anchored 11.6-kb dsRNA substrate. (B) Representative images (top) and kymographs (bottom) showing the Cy5-RIG-I binding to the free end of 11.6-kb dsRNA containing 5’-ppp. SYBR Gold stained 11.6-kb dsRNA is shown in green and Cy5-RIG-I (5 min incubation after injection) is shown in red. Merged images are generated by overlaying both channels. Positions of dsRNA are shown adjacent to the right of kymographs. Blue arrowheads indicate that Cy5-RIG-I molecules retracted with the dsRNA when buffer flow was turned off, suggesting protein binding to the dsRNA end. (C) Distribution of the positions for RIG-I binding on dsRNA (*n* = number of events). Diamonds represent individual events.

**Figure S4.**
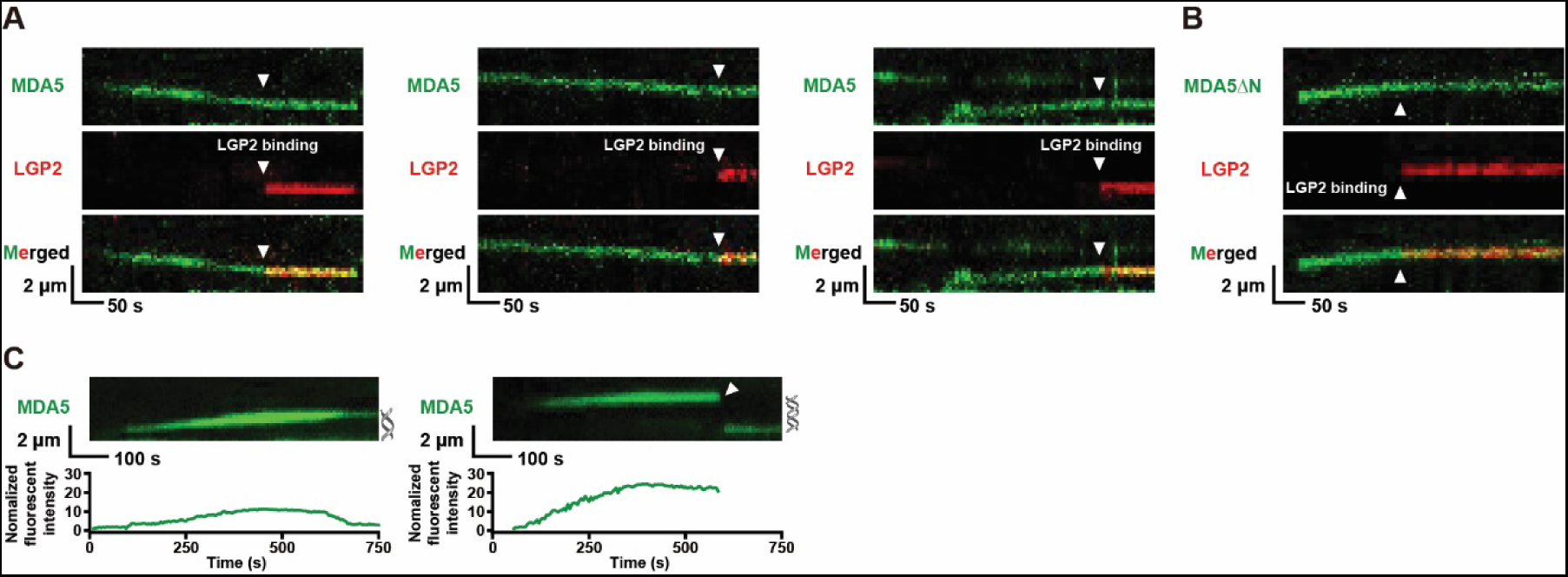
Kymographs of MDA5-LGP2 complexes and MDA5 foci. (A) Representative kymographs showing Cy5-LGP2 (3 nM) binding to Cy3-MDA5 motors (3 nM). Cy3-MDA5 are shown in green and Cy5-LGP2 are shown in red. Merged kymographs are generated by overlaying both channels. Arrowheads indicate the association of LGP2 with MDA5. (B) Representative kymographs showing a Cy5-LGP2 (0.5 nM) binding to a Cy3-MDA5ΔN motor (0.5 nM). Cy3-MDA5ΔN is shown in green and Cy5-LGP2 is shown in red. Merged kymograph is generated by overlaying both channels. Arrowheads indicate the association of LGP2 with MDA5ΔN. (C) Representative kymographs (top) and time-dependent fluorescent intensities (bottom) of Cy3-MDA5 foci. Arrowhead indicates the displacement of neutravidin from a dsRNA end. Positions of dsRNA are shown adjacent to the right of kymographs.

**Figure S5.**
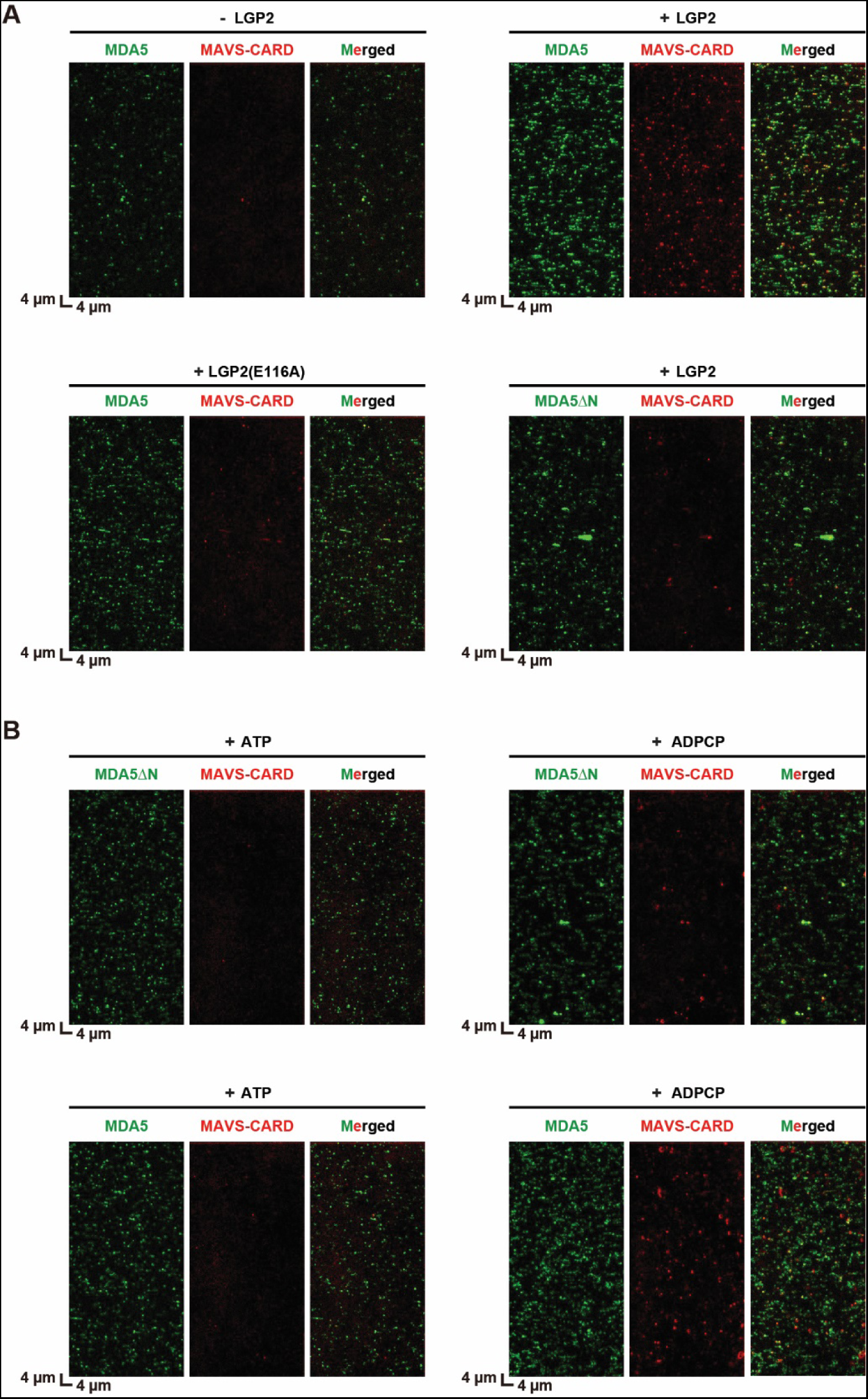
Images of MAVS-CARD recruitments by MDA5 foci. (A) Representative images showing the recruitments of AF647-MAVS-CARD (100 nM) by Cy3-MDA5 or Cy3-MDA5ΔN (20 nM) under various conditions. The absence or presence of LGP2/LGP2(E116A) (20 nM) is indicated above each set of images. (B) Representative images showing the recruitments of AF647-MAVS-CARD (100 nM) by Cy3-MDA5 or Cy3-MDA5ΔN (100 nM) under various conditions. The presence of ATP or ADPCP is indicated above each set of images.

**Figure S6.**
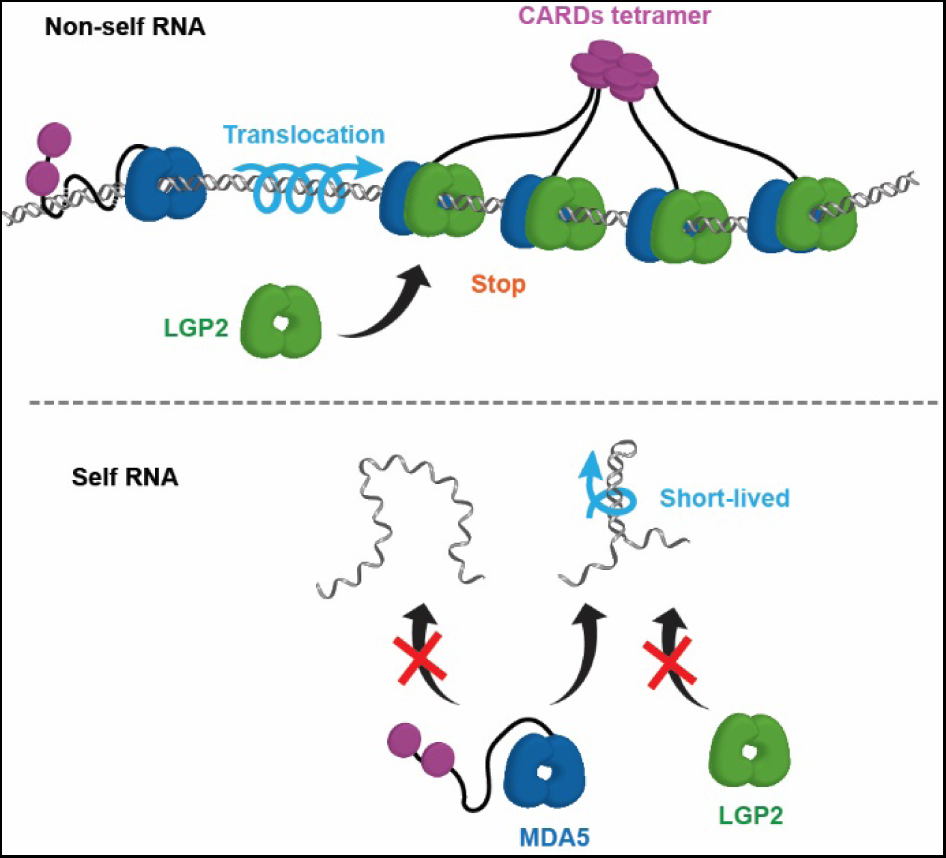
A translocation model for MDA5 signaling initiation. A proposed translocation model for MDA5 signaling. Top: MDA5 motors perform long-distance 1D movement on viral dsRNA to recruit LGP2, which further allows CARDs to assemble into functional tetramers to activate MAVS. During the rotation-coupled translocation of MDA5, the flexible linker between the CARDs and helicase domain wraps around the dsRNA backbone, resulting in restricted flexibility in CARDs arrangement. Bottom: MDA5 translocation on self RNA is likely to be short-lived and insufficient for LGP2 recruitment.

## SUPPLEMENTAL INFORMATION

Movie S1. Representative movie (20 frames per sec) of an MDA5 motor translocating on dsRNA, related to Figure 1. Two channels were merged. MDA5 is shown as green and dsRNA is shown as red.

Movie S2. Representative movie (20 frames per sec) of the formation of an MDA5-LGP2 complex on dsRNA, related to Figure 3. MDA5 is shown as green (top), LGP2 is shown as red (middle) and two channels were merged (bottom). Approximate position of dsRNA is shown as a white line. Co-localization between MDA5 and LGP2 is observed as yellow.

Movie S3. Representative movie (20 frames per sec) of MDA5 foci formation on dsRNA, related to Figure 4. Two channels were merged. MDA5 is shown as green and dsRNA is shown as red.

