## Supplementary Information for "Observation of Higher-Order Assemblies Controlled by Protein One-dimensional Movement"

### Supplementary Tables

### Table S1. Oligonucleotides Used in this study.

| Name | Sequences (5'-3') |
|---|---|
| MDA5 for1 | ATCACAGCAGCCTCCCTGAAACTGGAGGAGGGTCCATGTCGAATGGGTAT |
| MDA5 for2 | CAGAGAGGATCCATGCATCATCATCATCATCACAGCAGCCTC |
| MDA5 rev | AAGCTTGGTACCTAATCCTCATCACTAAATAAAC |
| MDA5(Q57E) for | GAGGCAGTTGAACTGCTGCTGAGCA |
| MDA5(Q57E) rev | CATGTTCCCGGAGGTGGCGACTGTC |
| MDA5ΔN for | ATGGGAAGTGATTCAGATGAAGAGA |
| MDA5ΔN rev | GGACCCTCCTCCAGTTTCAGGGAG |
| LGP2 for | CCCACCATCGGGCGCGGATCCATGGAGCTTCGGTCATACC |
| LGP2 rev1 | GGCTGCTACCGGTTTCCGGCAGGGATCCACCACCACCACCACCACCGTCCAGGGAGAGGT |
| LGP2 rev2 | CCTTTGAATTCCGCGCGCTTCGGACCGTTAATGATGATGATGATGATGGCTGCTACCGGTT |
| LGP2(K30A) for | GCCACCCGGGCGGCTGCTTATGTG |
| LGP2(K30A) rev | CCCGGCACCCGTGGGCAGCCAGA |
| LGP2(E116A) for | GCAGAGGAGCACGTGGAGCTCACT |
| LGP2(E116A) rev | CTCGGGGCTGGTCAGTGCCATCT |
| RIG-I for1 | TCATCACTCCCTGAAACTGGAGGAGGATCCATGACCACCGAGCAG |
| RIG-I for2 | GCTCACTAGTCGCGGCCGCATGCATCATCATCATCATCACTCATCACTCCCTGAA |
| RIG-I rev | GCATGCCTCGAGTCATTTGGACATTTCTGCTGGATC |
| MAVS-CARD for1 | AAGGTTGTAAAGATTCTGCCTTACCTGCCCTGCCT |
| MAVS-CARD for2 | AAGCTAGTTGATCTCGCGGACGAAG |
| MAVS-CARD rev1 | CACATTGCAAAAATTGCTGAAATTG |
| MAVS-CARD rev2 | ACAGCCCCTCAGTGCCGCAATGAAG |
| LGP2-Flag for | AAGCTTGCGGCCGCGAATTCAATGGAGCTTCGGTCATACCAATGGGA |
| LGP2-Flag rev | TATCAGATCTATCGAGAATTCGTCCAGGGAGAGGTCCGACAAGTTCTC |
| Myc-MDA5 for | GCCATGGAGGCCCGAATTCGGATGTCGAATGGGTATTCCACAGACGA |
| Myc-MDA5 rev | GAGAGATCTCGGTCGAGAATTCCTAATCCTCATCACTAAATAAACAG |

| | |
|---|---|
| RNA linker 1 | CCGGAUCGCUCGAGACGCAUUUGCAUCUAGAGGGCCCUAUUC-biotin |
| RNA linker 2 | GUACCGCGCUGAUAAGCCUGGUUGACGGAAGUGGCAAUUCUAGAGGGCCC UAUUC-biotin |
| ssRNA31 | UUUUUUGGGUUUUCCCAGUCACGACGUUGUA |
| MDA5 sgRNA1 for | CACCGTAGCGGAAATTCTCGTCTG |
| MDA5 sgRNA1 rev | AAACCAGACGAGAATTTCCGCTAC |
| MDA5 sgRNA2 for | CACCGGGTTGGACTCGGGAATTCG |
| MDA5 sgRNA2 rev | AAACCGAATTCCCGAGTCCAACCC |
| LGP2 sgRNA for | CACCGAGCTTCGGTCCTACCAAT |
| LGP2 sgRNA rev | AAACATTGGTAGGACCGAAGCTC |
| qPCR primer for IFNB for | CTTTCGAAGCCTTTGCTCTG |
| qPCR primer for IFNB rev | CAGGAGAGCAATTTGGAGGA |
| qPCR primer for ACTB for | CACTCTTCCAGCCTTCCTTC |
| qPCR primer for ACTB rev | TACAGGTCTTTGCGGATGTC |

### Table S2. Labeling efficiencies of proteins.

| Protein | Labeling Efficiencies |
|---|---|
| Cy3 labeled MDA5 | 50% |
| Cy3 labeled MDA5(Q57E) | 49% |
| Cy3 labeled MDA5ΔN | 30% |
| Cy3 labeled LGP2 | 55% |
| Cy5 labeled LGP2 | 39% |
| Cy3 labeled LGP2(K30A) | 51% |
| AF647 labeled MAVS-CARD | 49% |
| Cy5 labeled RIG-I | 18% |

**Movie S1.**

Representative movie (20 frames per sec) of an MDA5 motor translocating on dsRNA (Figure 1C, middle). Two channels were merged. MDA5 is shown as green and dsRNA is shown as red.

**Movie S2.**

Representative movie (20 frames per sec) of the formation of an MDA5-LGP2 complex on dsRNA (Figure 3A). MDA5 is shown as green (top), LGP2 is shown as red (middle) and two channels were merged (bottom). Approximate position of dsRNA is shown as a white line. Co-localization between MDA5 and LGP2 is observed as yellow.

**Movie S3.**

Representative movie (20 frames per sec) of MDA5 foci formation on dsRNA. Two channels were merged. MDA5 is shown as green and dsRNA is shown as red.
